# Lipid perturbation-activated IRE-1 modulates autophagy and lipolysis during endoplasmic reticulum stress

**DOI:** 10.1101/285379

**Authors:** Jhee Hong Koh, Lei Wang, Caroline Beaudoin-Chabot, Guillaume Thibault

## Abstract

Metabolic disorders such as obesity and nonalcoholic fatty liver disease (NAFLD) are emerging diseases that affect the global population. One facet of these disorders is attributed to the disturbance of membrane lipid composition. Perturbation of endoplasmic reticulum (ER) homeostasis through changes in membrane phospholipid composition results in activation of the unfolded protein response (UPR) and causes dramatic translational and transcriptional changes in the cell. To restore cellular homeostasis, the three highly conserved UPR transducers ATF6, IRE1, and PERK mediate cellular processes upon ER stress. The role of the UPR in proteotoxic stress caused by the accumulation of misfolded proteins is well understood but much less so under lipid perturbation-induced UPR (UPR^LP^). We found that genetically disrupted phosphatidylcholine synthesis in *C. elegans* causes, lipid perturbation, lipid droplet accumulation, and induced ER stress, all hallmarks of NAFLD. Transcriptional profiling of UPR^LP^ animals shows a unique subset of genes modulated in an UPR-dependent manner that is unaffected by proteotoxic stress (UPR^PT^). Among these, we identified autophagy genes *bec-1* and *lgg-1* and the lipid droplet-associated lipase *atgl-1* to be modulated by IRE-1. Considering the important role of lipid homeostasis and how its impairment contributes to the pathology of metabolic diseases, our data uncovers the indispensable role of a fully functional UPR program in regulating lipid homeostasis in the face of chronic ER stress and lipotoxicity.

## INTRODUCTION

Lipid content in a cell is tightly regulated to maintain cellular homeostasis and plays important roles in many physiological processes such as energy storage, signalling and contributes to membrane structure. A disruption in lipid homeostasis is associated with obesity and nonalcoholic fatty liver disease (NAFLD) (Puri et al., 2007, Tiniakos et al., 2010, Arendt et al., 2013, Doycheva et al., 2017). Lipid perturbation is often characterised by excessive accumulation of lipids in tissues such as liver, pancreas and adipose as a consequence of the inability to control detrimental processes such as lipotoxicity, dysfunctional unfolded protein response and apoptotic pathways that ultimately leads to a disease state (Hotamisligil and Erbay, 2008, Rinella and Sanyal, 2015). Altered hepatic phospholipids content occurs in mice model with nonalcoholic hepatosteatosis (NASH), a severe form of NAFLD (Li et al., 2006, Fu et al., 2011a). A decrease in the major phospholipid species phosphatidylcholine (PC) relative to phosphatidylethanolamine (PE) predicts survival and liver function after partial hepatectomy in mice (Ling et al., 2012). As both PC and PE are the two major phospholipids in the endoplasmic reticulum (ER) membrane, simultaneously blocking the two PC biosynthesis pathways from the ablation of the gene encoding for PE N-methyltransferase (PEMT) and the deprivation of dietary choline in mice leads to hepatic ER stress which correlate with steatosis progression (Li et al., 2005) (Fig. 1A). These observations link lipid perturbation, ER stress, and metabolic diseases.

**Figure 1.**
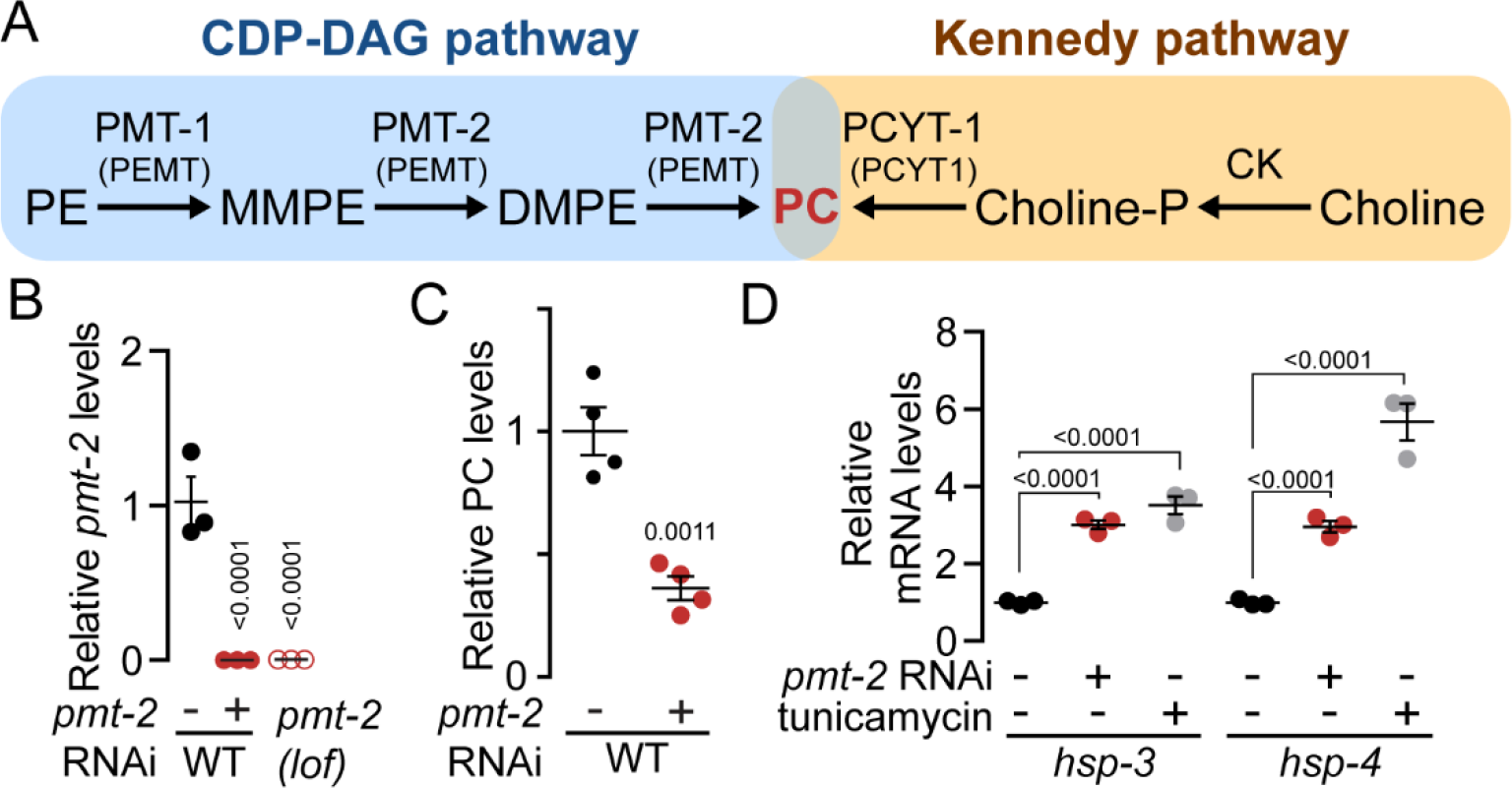
*pmt-2* silencing is sufficient to activate the UPR from lipid perturbation. (A) Metabolic pathways for the synthesis of phosphatidylcholine in *C. elegans*. Human orthologues are highlighted in brackets. PE, phosphatidylethanolamine; MMPE, monomethyl-phosphatidylethanolamine; DMPE, dimethyl-phosphatidylethanolamine; PMT-1/2, phosphatidylethanolamine N-methyltransferase 1/2; PEMT, phosphatidylethanolamine N-methyltransferase; PCYT-1, choline-phosphate cytidylyltransferase 1; CK, choline kinase. (B) qPCR results comparing expression of *pmt-2* in *pmt-2(RNAi)* and in *pmt-2(lof)* animals (C) Comparison of PC levels in WT and *pmt-2(RNAi)* animals by GC-FID. (D) qPCR comparing expression of UPR marker genes *hsp-3* and *hsp-4* in *pmt-2(RNAi)* and WT worms treated with 25 µg/ml tunicamycin for 4 h.

The ER is the hub of protein folding targeted to the secretory pathway (Schroder and Kaufman, 2005, Braakman and Bulleid, 2011). In addition to its role in protein homeostasis, the ER is the site of lipid metabolism providing the majority of membrane lipids throughout the cell (Lagace and Ridgway, 2013, Wu et al., 2014). Long term disruption of lipid homeostasis triggers chronic ER stress which is associated with metabolic diseases (Fu et al., 2011a, Han and Kaufman, 2016). To counter ER stress, eukaryotes have evolved transcriptional and translational regulatory pathways termed the unfolded protein response (UPR) to ease stress and maintain ER functionality (Cox and Walter, 1996, Schroder and Kaufman, 2005, Shen et al., 2004). The UPR, which is highly conserved from yeast to humans, exerts homeostatic control by sensing stress signals arising from the ER lumen through misfolded protein accumulation or from membrane lipid perturbation (Walter and Ron, 2011, Volmer et al., 2013). The signal is then transduced in the nucleus by upregulating a subset of genes to remodel various cellular pathways to manage the existing stress and maintain viability. Failure of UPR intervention to reach ER homeostasis may result in cell death through apoptosis (Oyadomari et al., 2002, Wu et al., 2014, Mota et al., 2016). Therefore, maintenance of ER homeostasis is essential for cellular functions and in the survival of eukaryotes.

The UPR consists of three conserved ER-localised stress sensing transmembrane transducers namely ATF6, IRE1 and PERK. Upon ER stress, each sensor activates their downstream effectors resulting in the combination of general translational shutdown and the upregulation of a subset of genes to restore cellular homeostasis (Walter and Ron, 2011, Wu et al., 2014). In the event of an acute response to excessive ER stress, the UPR is known to elicit cell death response mediated through IRE1 and PERK (Harding et al., 2003, Novoa et al., 2001). In addition, chronic ER stress characterised by the continual burden on the ER function has surprisingly resulted to an adaptative response by the UPR (Rutkowski et al., 2006, Kim et al., 2017). Contrary to the apoptotic response observed in acute ER stress, adaptation to chronic ER stress is characterised by resistance to apoptosis resulting from differential expression of pro-survival signals and stabilisation of UPR components (Rubio et al., 2011).

Previously, we have demonstrated that changes in membrane lipid composition through the ablation of the *de novo* PC biosynthesis gene *OPI3* (*PEMT* orthologue) activate the essential intervention of the UPR to remodel the protein homeostasis network in budding yeast (Thibault et al., 2012, Ng et al., 2017). The UPR can be directly activated from lipid perturbation at the ER membrane independently from misfolded proteins accumulating in the ER lumen (Promlek et al., 2011, Volmer et al., 2013, Ho et al., 2018). Similar to the NAFLD mice model lacking *PEMT* gene (Li et al., 2006, Li et al., 2005) and our yeast model missing *OPI3* gene (Thibault et al., 2012, Ng et al., 2017), NAFLD models have been developed in *C. elegans* to directly prevent PC synthesis from the loss-of-function *(lof)* of *PEMT* orthologues *pmt-1* and *pmt-2* or the upstream precursor S-adenosylmethionine synthetase *sams-1* (Brendza et al., 2007, Li et al., 2011, Walker et al., 2011, Ding et al., 2015) (Fig. 1A). Generally, perturbing PC level affects the abundance and size of lipid droplets, serving as a compensatory response to decreased PC that results in channelling neutral lipids into lipid droplets (Guo et al., 2008, Li et al., 2011, Walker et al., 2011). Lipid droplets are intracellular organelles that store neutral lipids, triacylglycerol and sterol, with increasing emergence of its role linked to cellular functions (Gross and Silver, 2014, Martinez-Lopez and Singh, 2015, Welte, 2015, Hashemi and Goodman, 2015, Nguyen et al., 2017).

In this study, we characterised the UPR function during lipid perturbation-induced chronic ER stress. We hypothesised that lipid perturbation may elicit chronic ER stress that is distinctly different from acute ER stress. In *C. elegans*, we silenced *pmt-2* to attenuate PC synthesis in the UPR mutant animals; *atf-6(lof)*, *ire-1(lof)* and *pek-1(lof)*. As expected, genetic attenuation of *pmt-2* results in the perturbation of lipid homeostasis with large decrease in PC levels which correlate with UPR activation. Although conventionally seen as a linear response from ER stress, our findings demonstrate a strikingly different outcome of the UPR programme when activated from lipid perturbation-induced (UPR^LP^) as opposed to proteotoxic-induced (UPR^PT^) ER stress. Interestingly, the data indicate that the ER stress sensor IRE-1 regulates the autophagy pathways and lipolysis suggesting a reciprocal relationship between lipid homeostasis and autophagy as mediated by the UPR to prevent excessive lipid droplet accumulation.

## RESULTS

### The attenuation of *pmt-2* activates the UPR by reducing total phosphatidylcholine

*PMT-1* and *PMT-2* are both required for phospholipid synthesis of PC from PE in *C. elegans* (Palavalli et al., 2006, Brendza et al., 2007, Li et al., 2011) (Fig. 1A). In the absence of choline, both genes are essential for the development of *C. elegans* and silencing either genes from stage-one (L1) larva leads to sterility (Brendza et al., 2007). PC cannot be obtained from standard laboratory worm diet as the conventionally used *E. coli* strains OP50 and HT115(DE3) lack PC (Morein et al., 1996, Oursel et al., 2007). Thus, PC levels can be altered by genetically manipulating *pmt-1* or *pmt-2* to induce lipid perturbation (LP). To better understand the role of the UPR during membrane LP, we subjected synchronised L1 larval stage worms to *pmt-2* RNA interference (RNAi) for 48 h. Two-day RNAi feeding was sufficient to decrease *pmt-2* mRNA in wild-type (WT) animals close to the background signal of *pmt-2(lof)* (Fig. 1B). As previously reported, *pmt-2(RNAi)* animals showed a developmental defect characterised by reduced body size which can be rescued through choline supplementation (Fig. S1A,B) (Palavalli et al., 2006). PC is synthesised from choline through the Kennedy pathway in *pmt-2(RNAi)* animals (Fig. 1A). Supplementing *pmt-2(RNAi)* animals with 30 mM choline was sufficient to prevent UPR activation while the growth defect was alleviated by 60 mM choline supplementation (Fig. S1C). To further characterise *pmt-2(RNAi)* worms, PC was separated from total lipid extract by thin layer chromatography (TLC) followed by transesterification to derive fatty acid methyl esters (FAMEs) from PC and quantified by gas chromatography with flame ionisation detector (GC-FID). Only 36% of PC remained in *pmt-2(RNAi)* worms compared to vector control (Fig. 1C). As LP can lead to ER stress, we measured the transcriptional levels of UPR-induced ER-resident molecular chaperone Hsp70s (Urano et al., 2002). The mRNA level for both human BiP orthologues, *hsp-3* and *hsp-4*, were upregulated transcriptionally when compared to WT worms (Fig. 1D). The mRNA level of *hsp-3* in *pmt-2(RNAi)* was similar to WT worms incubated 4 h with the strong UPR inducer tunicamycin (Tm). Tm blocks all N-glycosylation of proteins leading to a severe accumulation of unfolded proteins in the ER (Ericson et al., 1977). In contrast, mRNA level of *hsp-4* from Tm treatment was remarkably higher compared to LP. This result suggests that *hsp-4* might be modulated differently from the UPR program when activated from LP (UPR^LP^) or proteotoxic (UPR^PT^) stress. As the UPR is activated from low PC, *C. elegans* subjected to *pmt-2* RNAi is a relevant model of NAFLD where altered PC/PE ratios and UPR activation are interconnected in the liver (Ozcan et al., 2004, Fu et al., 2011b, Thibault et al., 2012, Ng et al., 2017).

### Attenuated phosphatidylcholine synthesis leads to lipid droplet accumulation

To investigate the regulatory role of the three ER stress transducers during LP (Shen et al., 2005), *atf-6(lof), ire-1(lof)* and *pek-1(lof)* mutant worms were subjected to *pmt-2* RNAi as described above (Fig. S2A). The ablation of two or three UPR branches is not possible as any combinations are lethal (Shen et al., 2005). As expected from *pmt-2* RNAi, reduction of PC level across the three UPR mutant worms is comparable to WT worms (Fig. 2A, S2B). To quantify fatty acids (FAs) composition of PC and its precursor PE during lipid perturbation, both phospholipids were separated from total lipid extract by TLC and quantified by GC-FID as described above. As expected, PC depletion in UPR mutants and WT worms caused significant reduction of FAs derived from PC at the exception of margaric acid (C17:0), α-linolenic acid (C18:3n3) and eicosadienoic acid (C20:2), indicating a general disturbance in the PC metabolic pathways during lipid perturbation (Fig. 2B). However, levels of FAs derived from PE were largely unmodified across the strains as *pmt-2* RNAi leads to the accumulation of the phospholipid intermediate monomethyl-phosphatidylethanolamine (MMPE) (Fig. 1A). As a reduction in PC level leads to lipid droplet (LD) accumulation in eukaryotes (Li et al., 2011, Walker et al., 2011, Thibault et al., 2012, Horl et al., 2011), Sudan Back staining of fixed *pmt-2(RNAi)* worms was carried out to visualise lipid droplets by Nomarski optics microscopy (Fig. 2C, S3A). We observed increased body fat in WT, *atf-6(lof)*, *ire-1(lof)*, and *pek-1(lof)* worms treated with *pmt-2* RNAi, suggesting increased lipid accumulation when PC is reduced. Increased fat accumulation as a consequence of PC perturbation has been demonstrated through silencing of *sams-1* and *pmt-1* RNAi (Ding et al., 2015).

**Figure 2.**
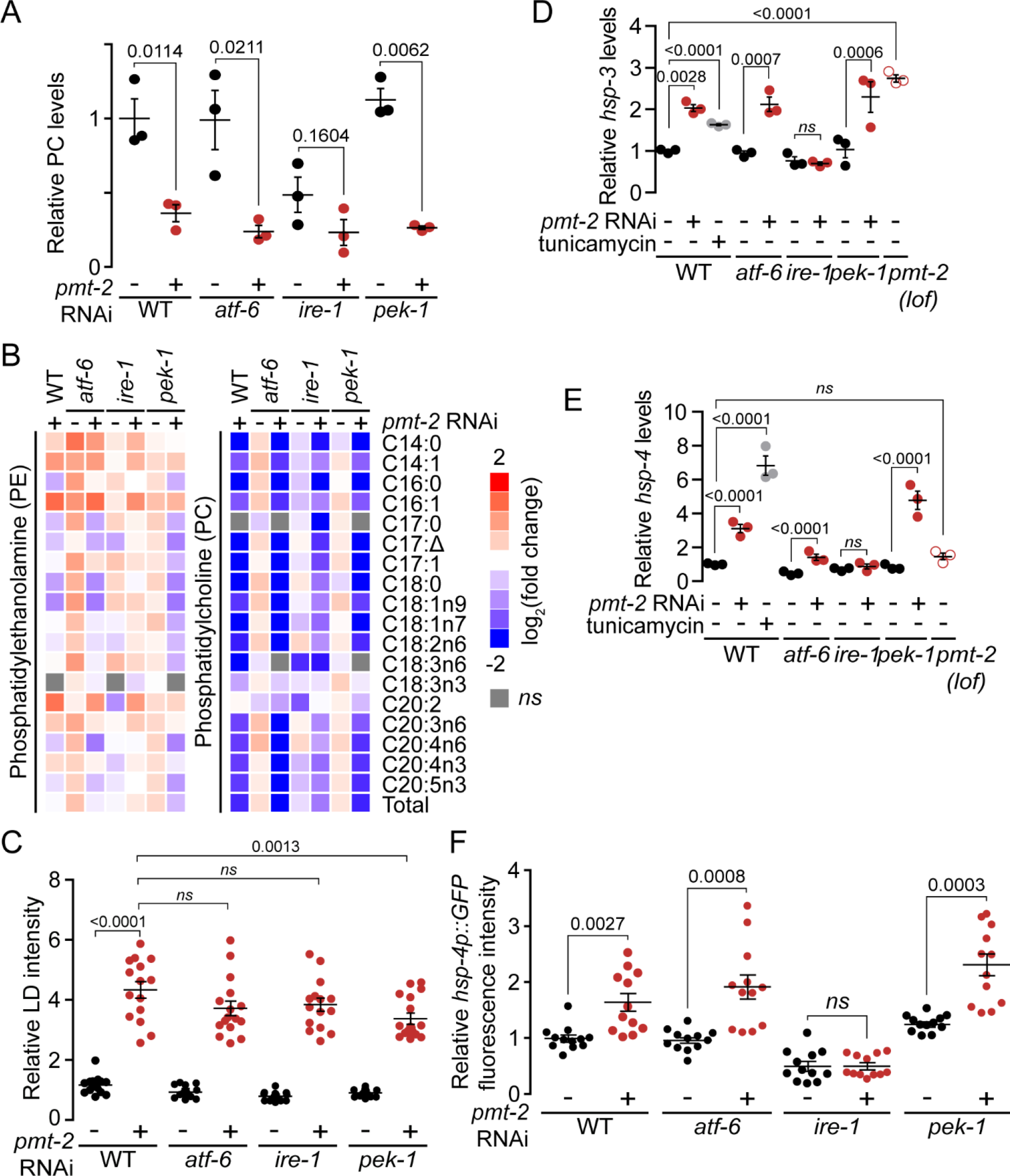
UPR^LP^ leads to the accumulation of lipid droplets. (A) Comparison of PC levels in WT, *atf-6(lof)*, *ire-1(lof)* and *pek-1(lof)* animals treated with *pmt-2* RNAi by GC-FID. (B) Heat maps reflecting all significant PE- and PC-derived fatty acids changes to untreated WT. WT and mutant worms were treated as in A. (C) Lipid droplet (LD) quantified from Sudan Black B staining fluorescence images of WT and mutant animals treated as in A. n > 16. (D, E) qPCR comparing expression of UPR marker genes *hsp-3* and *hsp-4* in WT and mutant animals treated as in A and WT worms treated with 25 µg/ml tunicamycin for 4 h. (F) Quantification of UPR reporter gene *hsp-4::GFP* of fluorescence images in WT and mutant animals treated as in A. n = 60. *ns* = non-significant.

To ensure that *pmt-2* RNAi treatment is sufficient to induce ER stress in the UPR mutants, we monitored the mRNA expression of two downstream target genes of IRE-1, *hsp-3* and *hsp-4*, by quantitative RT-PCR. As expected, *hsp-3* levels were significantly increased upon *pmt-2* RNAi and tunicamycin treatments in WT and mutant strains with the exception of *ire-1(lof)* (Fig. 2D). Similarly, this ER-resident chaperone hsp70 was significantly increased in *pmt-2(lof)*. In contrast, *hsp-4* levels were not significantly different in WT, *atf-6(lof)*, and *ire-1(lof)* treated with *pmt-2* RNAi as well as in *pmt-2(lof)* when compared to untreated WT animals (Fig. 2E). *hsp-4* levels were only significantly increased in *pek-1(lof):pmt-2(RNAi)* mutant and WT treated with tunicamycin. IRE1 compensates for the lack of PERK resulting in higher BiP (*hsp-4* orthologue) upregulation upon ER stress (Yamaguchi et al., 2008). It should be noted that *hsp-4* was found to be significantly upregulated in *pmt-2(RNAi)* worms (Fig. 1D). However, the fold change of *hsp-4* expression is similar in both experiments and mild when compared to tunicamycin treatment (Fig. 1D, 2E). The activation of the UPR from LP was further investigated *in vivo* by using the *hsp-4p::GFP* reporter animal (Calfon et al., 2002). Fluorescence was significantly increased upon *pmt-2* RNAi treatment in WT, *atf-6(lof)* and *pek-1(lof)* worms but not in *ire-1(lof)* animal (Fig. 2F and S3B). This discrepancy from the qPCR results might be due to the accumulation of stable GFP from mild but prolonged ER stress while *hsp-4* mRNA turnover can be rapid. Overall, genetic attenuation of PC synthesis activates the UPR in which the three UPR branches are required to prevent the accumulation of LDs.

### UPR^LP^ upregulates a distinct subset of genes from UPR^PT^

Several studies suggest that the essential role of the UPR in maintaining metabolic and lipid homeostasis is highly conserved across species (for reviews, see (Volmer and Ron, 2015, Han and Kaufman, 2016)). Recently, a novel ER stress sensing mechanism was proposed in which the UPR is activated from lipid perturbation independently by the accumulation of unfolded protein in the ER (Volmer et al., 2013). Thus, ER stress triggered from proteotoxic stress or lipid perturbation might differentially modulate the UPR to reach cellular homeostasis. To answer this question, DNA microarray analysis was performed using RNA extracted from WT, *atf-6(lof)*, *ire-1(lof)* and *pek-1(lof)* animals treated with *pmt-2* RNAi (GEO: GSE99763). WT worms incubated 4 h with tunicamycin were included in the analysis to identify genes modulated by the UPR^PT^ and subsequently uncouple those specifically modulated by the UPR^LP^ programme. To validate the quality of microarray data, qPCR was performed on a gene subset (Fig. S4). Overall, 2603 and 1745 genes were upregulated and downregulated, respectively, in *pmt-2(RNAi)* animals compared to WT (Fig. 3A, Supplementary file 1). In addition, tunicamycin-treated WT worms showed upregulation of 1258 genes and downregulation of 1473 others. Only 492 and 420 genes were similarly upregulated and downregulated, respectively, from both UPR^LP^ and UPR^PT^ suggesting these groups of genes are commonly modulated from the UPR regardless of the source of ER stress.

**Figure 3.**
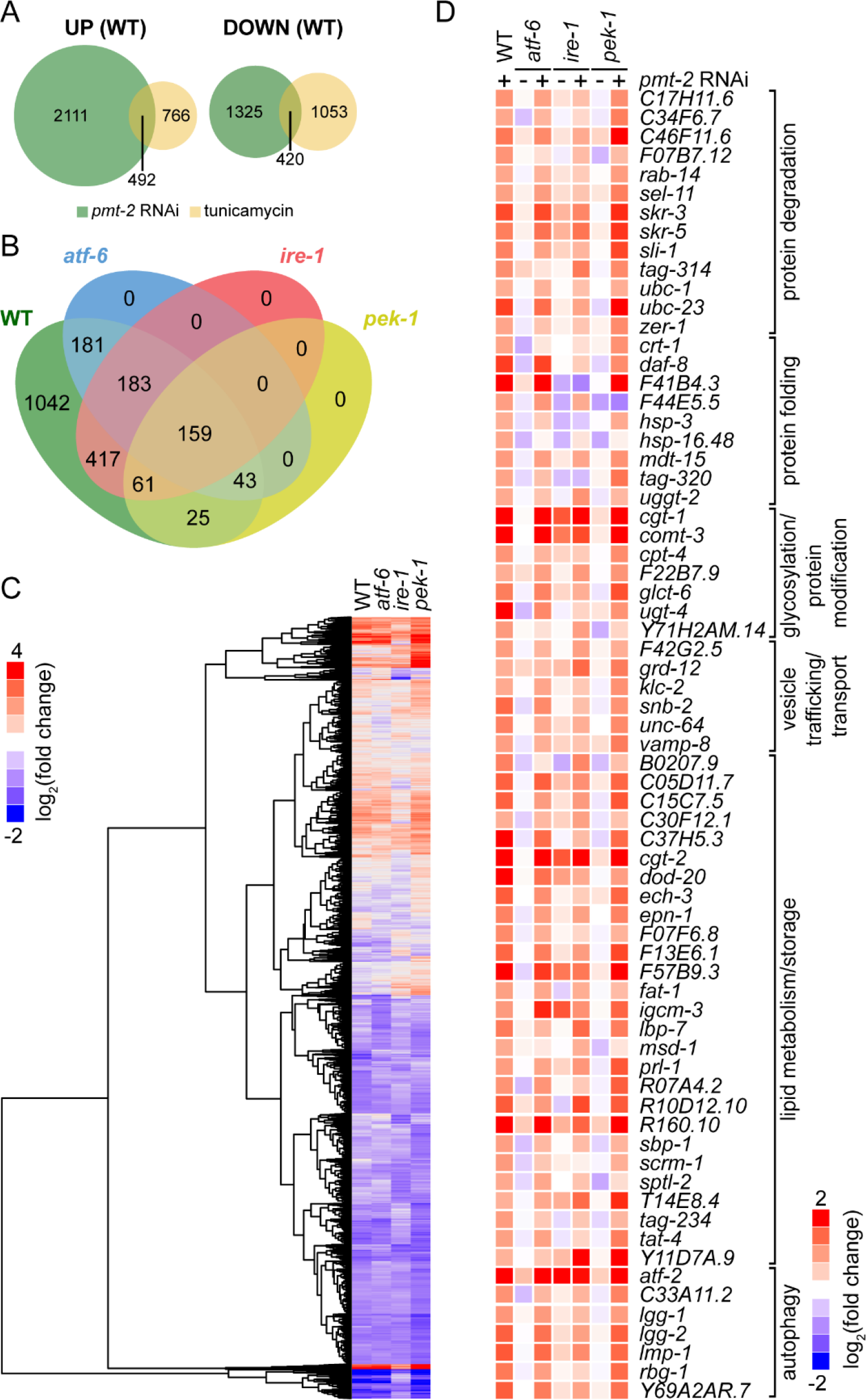
UPR^LP^ and UPR^PT^ lead to different transcriptomic outcomes. (A) Venn diagram representation of upregulated (UP) and downregulated (DOWN) genes at a minimum of 1.5 fold in *pmt-2(RNAi)* and WT worms treated with 25 µg/ml tunicamycin for 4 h compared to untreated WT worms. (B) Four-way Venn diagram depicts transcriptional targets of WT, *atf-6(lof), ire-1(lof)* and *pek-1(lof)* worms upregulated during lipid perturbation and excluding genes commonly upregulated from *pmt-2* RNAi and tunicamycin treatments. Fold-change > 1.5, ANOVA *P* values < 0.05. (C) Hierarchical clustering of 8045 genes with significant changes based on Pearson correlation coefficients. Gene expression of WT, *atf-6(lof), ire-1(lof), pek-1(lof)* worms treated with *pmt-2* RNAi were compared against their respective untreated worms (empty vector). (D) Heat map of previously reported UPR-regulated genes of worms treated as in C.

To explore how the UPR elicits a differential stress response during lipid perturbation from that under proteotoxic stress, we filtered the 2111 gene candidates that were upregulated only from UPR^LP^ in *pmt-2(RNAi)* animals and excluding genes upregulated from UPR^PT^ in tunicamycin-treated animals. Furthermore, genes that were not upregulated in at least one of the three UPR mutants, subjected to *pmt-2* RNAi, are considered to be modulated by the UPR in a LP-dependent manner. From this analysis, 1069 and 1042 genes were upregulated in a UPR^LP^-dependent and independent manner, respectively (Fig. 3B, Supplementary file 2). We identified 181, 417, and 25 genes that are specifically upregulated from ATF-6, IRE-1, and PEK-1 branches, respectively, while 446 genes are modulated from at least two of the three UPR branches, suggesting compensatory roles of one or more UPR transducers in the absence of others. In addition, we grouped genes with at least 1.5-fold change by hierarchical clustering (Fig. 3C, Supplementary file 3). This allowed us to visualise genes that were similarly regulated throughout the array from UPR^LP^. Manual inspection of our array data demonstrates that the upregulation of known UPR target genes are in agreement with previous reports (Fig. 3D) (Travers et al., 2000, Shen et al., 2005, Thibault et al., 2011).

To further understand the role of UPR^LP^ programme, we performed functional annotation for the 1069, 181, 417, and 25 upregulated genes identified from *pmt-2(RNAi)*, *atf-6(lof)*; *pmt-2(RNAi)*, *ire-1(lof)*; *pmt-2(RNAi)*, and *pek-1(lof)*; *pmt-2(RNAi)* animals, respectively, using the gene ontology tool DAVID (Supplementary file 4) (Huang et al., 2007). Immune regulatory genes were found to be enriched in the upregulated categories of WT and *ire-1* animals (Fig. 4A, C). This is in agreement with a previous report where innate immunity was found to be modulated by *sams-1*, the methyl donor to *pmt-1* and *pmt-2* (Ding et al., 2015). Our data suggest that innate immune response is positively and negatively regulated by IRE-1 and PEK-1, respectively (Fig. 4C-D). As expected, GO terms related to ER stress were found to be enriched in the upregulated categories of WT and mutant animals when PC is depleted (Fig. 4A-D) (26321661). We also identified an enrichment of downregulated genes in WT related to translational initiation factor including *eif-1.A, eif-3.C* and *eif-3.E,* a characteristic of the UPR (Ling et al., 2009, Long et al., 2002).

**Figure 4.**
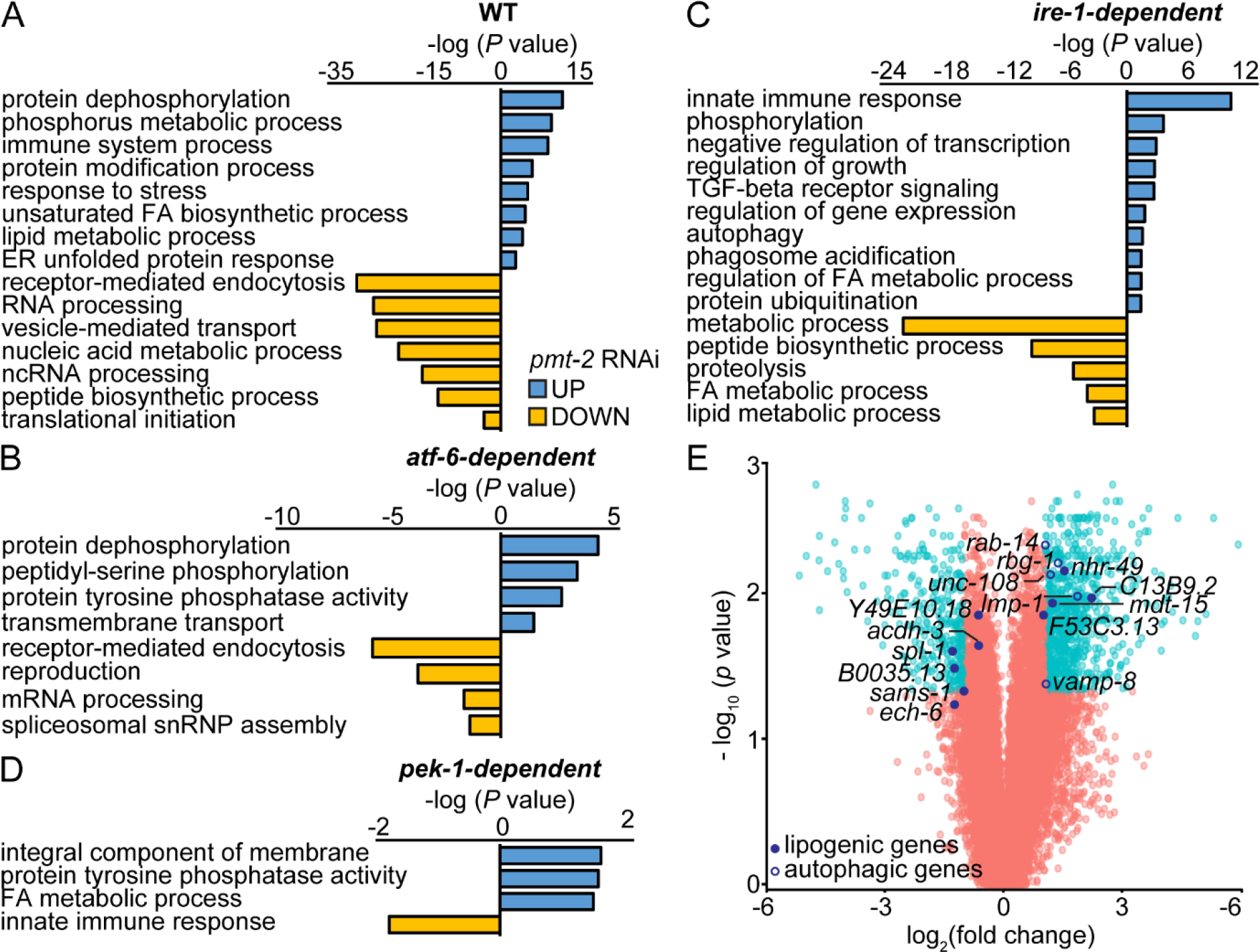
Depletion of PC increases autophagy and lipid metabolism activity in an IRE-1-dependent manner. (A-D) Bar plot of the GO analysis of genes upregulated (blue) and downregulated (yellow) in WT (A), *atf-6(lof)* (B)*, ire-1(lof)* (C) and *pek-1(lof)* (D) subjected to *pmt-2* RNAi and compared to their respective untreated strains (empty vector). Genes are highlighted in yellow in Supplementary file 4. (E) Volcano plot depicts gene expression of *pmt-2(RNAi)* compared to WT animals. Genes that are upregulated or downregulated by more than two fold binary logarithm and with false-discovery rate (FDR) < 0.05 are labelled in light blue circles. Lipogenic (closed circle) and autophagic (opened circle) genes regulated in an IRE-1-dependent manner are shown in dark blue.

Protein tyrosine phosphatase activity is significantly regulated by ATF-6 (GO ID: 0035335, n = 6, Benjamini *P* values 0.047) (Fig. 4B). This class of genes is activated in response to ER stress (Agouni et al., 2011). Transcriptional regulation (GO ID: 0006355, n = 49, Benjamini *P* values 3.3E-09) is enriched in IRE-1 and supports the evidence that it alleviates ER stress by widespread transcriptional modification process (Fig. 4C) (Ng et al., 2000). In addition, genes involved in lipid and fatty acid processes were also enriched by IRE-1 and PEK-1 (Fig. 4C-D). Lastly, protein tyrosine phosphatase activity (GO ID: 0004725, n = 4, Benjamini *P* values = 0.039) is enriched in PEK-1, suggesting its involvement in protein modifications and cell signalling cascade during lipid perturbation (Fig. 4D) (Bettaieb et al., 2012). Interestingly, we observed upregulation of autophagy-related processes that are IRE-1 dependent (Fig. 4 C,E). Crosstalk between the UPR and autophagy in the context of protein disaggregation has been well documented (for review see (Mizushima and Komatsu, 2011)). More recently, catabolic process of triacylglycerol (TG) breakdown to generate free fatty acids from lipid droplets through autophagy was proposed to be a necessary mechanism to maintain ER homeostasis in yeast (Velazquez et al., 2016). Thus, breakdown of TG to prevent acute ER stress might be mediated in a conserved IRE-1-dependent manner from the UPR^LP^ programme.

### IRE-1/XBP-1 axis regulates lipophagy during UPR^LP^

To investigate if the UPR^LP^ mediates TG hydrolysis through autophagy, we carried out an RNAi screen of autophagy-related genes. For the purpose of the screening, WT worms were first subjected to *pmt-2* RNAi for 36 h to induce UPR^LP^ followed by feeding of bacteria expressing autophagy-related RNAi genes for 5 days (Fig. 5A). Phenotypes following RNAi treatment were classified as 0 (little difference in growth and brood size), 1 (smaller brood size, sick), and 2 (sterile, very sick) compared to vector control. The screen was carried out twice and scores were designated for both WT animals pre-treated with vector and *pmt-2* RNAi. A clone was classified as positive if the difference of the sum of *pmt-2* RNAi treatment to vector treatment is ≥ 3 (Supplementary file 5). From the 40 RNAi clones, 9 were found to induce development defects upon RNAi treatment compared to vector control (Fig. S5). As positive controls, we incorporated *ero-1* and *pmt-2* RNAi as they both lead to developmental defect and sterility. The screen revealed that some autophagy-related genes are required during UPR^LP^ and the absence of autophagy contributes to observed detrimental phenotypes. Identified genes includes *atg-7* (autophagosome conjugation), *atg-13* (autophagosome formation), *bec-1* (vesicle nucleation), and *wdfy-3* (autophagy adaptor) (Wang et al., 2018, Jia et al., 2007, Takacs-Vellai et al., 2005, Melendez et al., 2003). Additionally, we included potential autophagy-related genes that are less characterised and prompt further investigation, these include *rsks-1*, *sepa-1*, and *trpp-8*.

**Figure 5.**
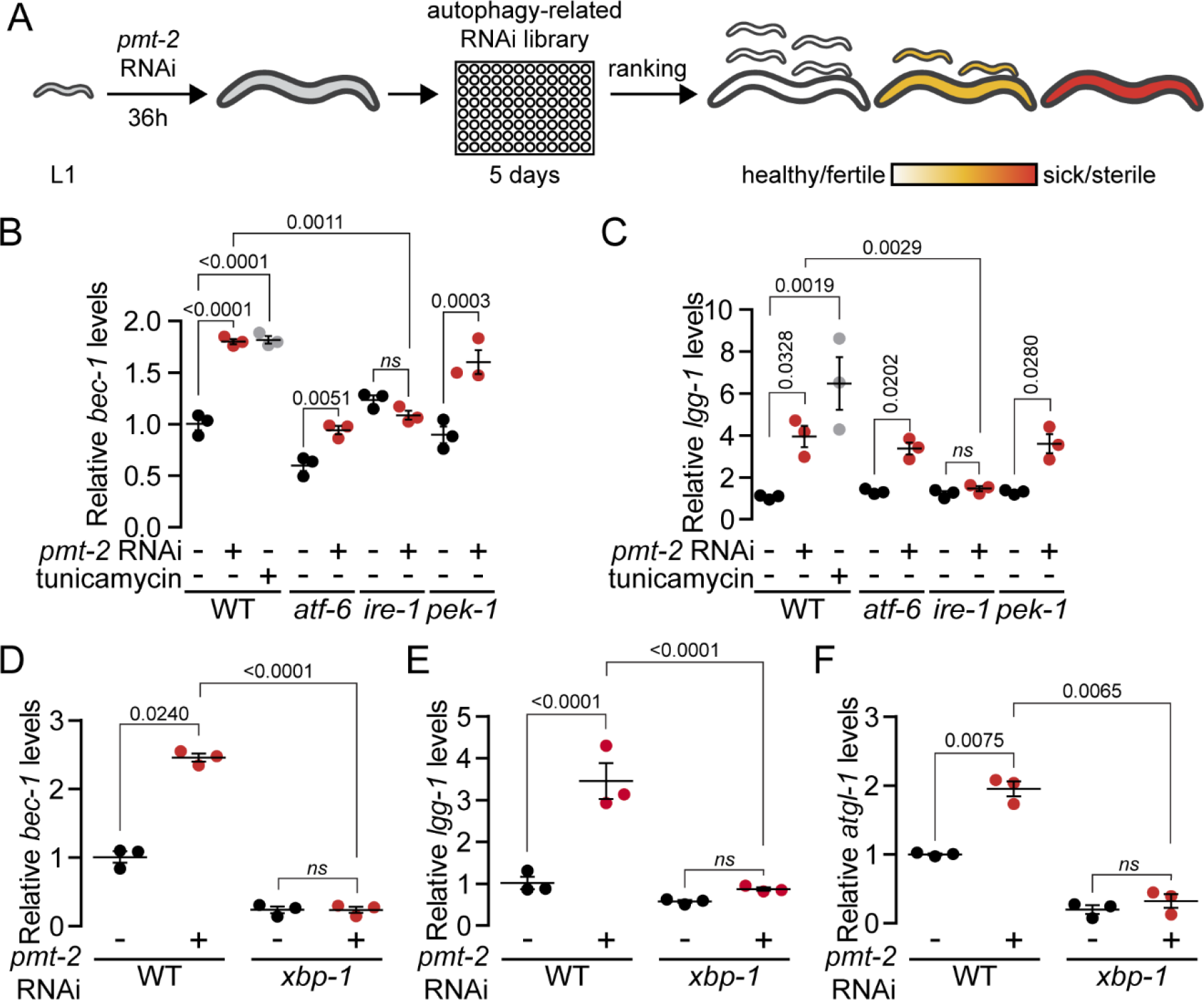
Autophagy genes *bec-1* and *lgg-1* are upregulated by IRE1/XBP1 axis upon UPR^LP^. (A) Schematic representation of the RNAi screening to identify potential autophagy genes that participate in fatty acid remodelling. WT worms were treated with *pmt-2* RNAi for 36 h and subsequently transferred to 96-well plates containing RNAi of autophagy-related target genes. The worms were scored for growth defects, sterility, mobility and embryonic lethality. qPCR comparing expression of *bec-1* (B) and *lgg-1* (C) in WT, *atf-6(lof), ire-1(lof)* and *pek-1(lof)* worms. qPCR comparing expression of *bec-1* (D), *lgg-1* (E) and *atgl-1(lof)* in WT and *xbp-1(lof)* mutants treated with *pmt-2* RNAi. ns, non-significant.

To better understand the crosstalk between UPR^LP^ and autophagy, we focused on the autophagy regulatory gene *bec-1* identified from the RNAi screening in addition of autophagy gene *lgg-1*. Both *bec-1* and *lgg-1* genes were found to be significantly upregulated in WT, *atf-6(lof)* and *pek-1(lof)* but not in *ire-1(lof)* from *pmt-2* RNAi treatment compared to control (Fig. 5B,C). These results suggest that IRE-1 is the only regulator of *bec-1* and *lgg-1* from the UPR^LP^ programme. We validated that *bec-1* and *lgg-1* are both upregulated from tunicamycin as previously reported (Ogata et al., 2006). Once IRE-1 is activated, it splices *xbp-1* mRNA through unconventional splicing resulting in the translation of the transcription factor XBP-1 which regulates downstream signalling cascade (Walter and Ron, 2011, Ho et al., 2018). XBP-1 is essential to upregulate *hsp-3* during UPR^LD^ (Fig. S5A). Upon prolonged ER stress, IRE-1 cleaves mRNAs to relieve protein load through its regulated IRE1-dependent decay (RIDD) activity (Hollien et al., 2009). To narrow down the role of IRE-1 in modulating autophagy, we monitored *bec-1 and lgg-1* mRNA levels in *xbp-1(lof)* treated with *pmt-2* RNAi. The upregulation of *bec-1* and *lgg-1* genes were abolished in *xbp-1(lof)* suggesting that both genes are regulated by XBP-1 during UPR^LP^ (Fig. 5D, E). As expected, *hsp-3* was not upregulated in *xbp-1(lof); pmt-2(RNAi)* (Fig. 6). Therefore, these results demonstrate that autophagy is regulated through the IRE-1/XBP-1 axis specifically from the UPR^LP^ programme.

**Figure 6.**
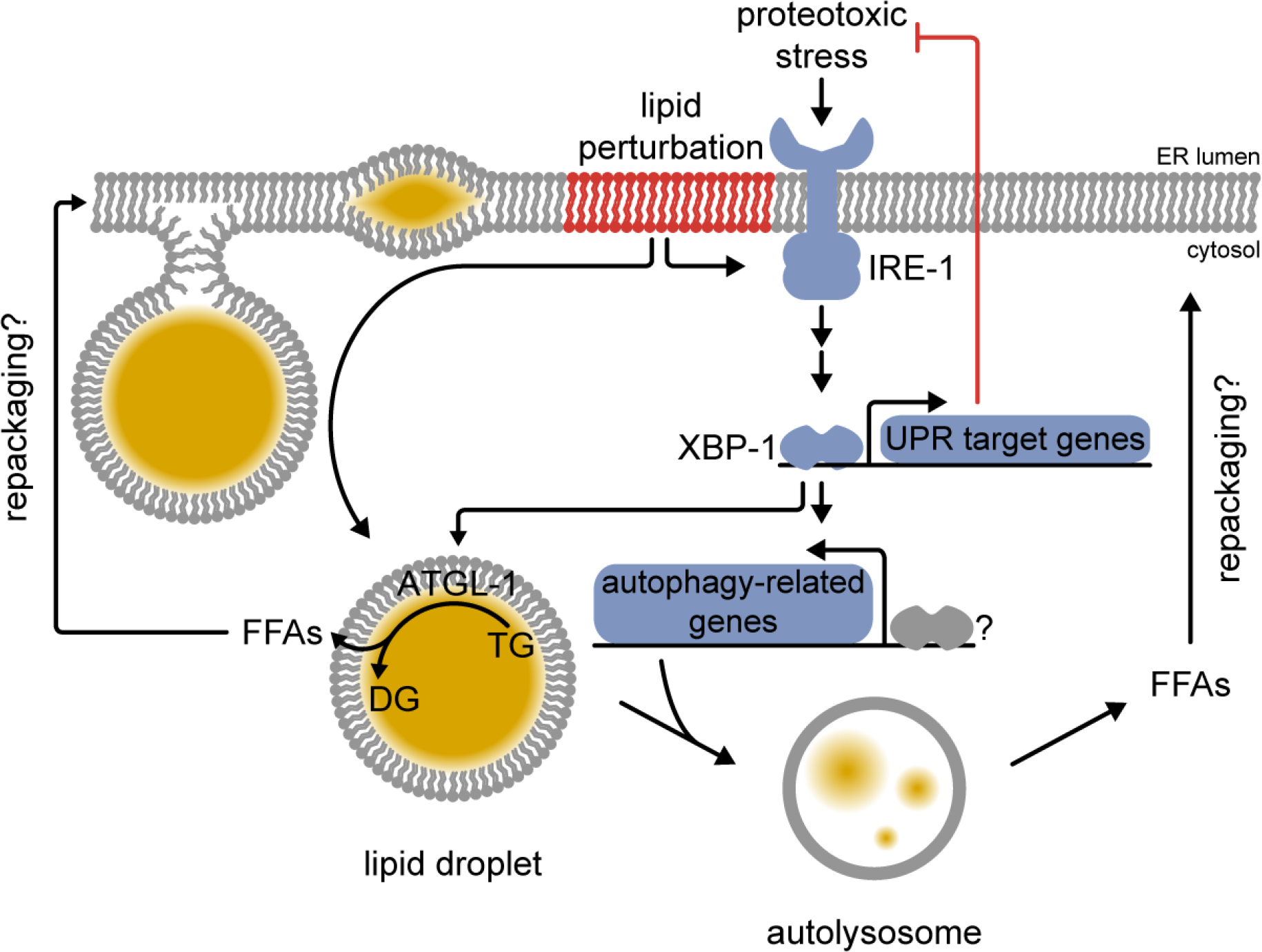
The UPR^LP^ maintains lipid homeostasis by selective autophagy. Proteotoxic-induced ER stress activates a subset of UPR target genes to restore ER homeostasis. In addition, lipid perturbation-induced ER stress induces the breakdown of excessive lipid droplets by autophagy which is regulated by the IRE-1/XBP-1 axis. This rechannelling of lipids into free fatty acids (FFAs) might be required to rapidly rebuild deficient phospholipids in an attempt of restoring lipid homeostasis at the ER. However, in the context of PC deficiency-induced NAFLD, this process will fail to restore lipid homeostasis leading to prolonged ER stress. DG, diacylglycerol; TG, triacylglycerol.

Turnover of lipid droplets is known to be mediated by cytosolic lipase ATGL-1 in *C. elegans*. ATGL-1 initiates TG hydrolysis to diacylglycerol (DG) and free fatty acid (FFA). Recently, studies have revealed the ATGL-1 modulate lipid breakdown through autophagy. In mammalian cells, ATGL (ATGL-1 orthologue) promotes lipophagy under the control of the transcription factor SIRT1 (Sathyanarayan et al., 2017). Lipid droplets turnover can be regulated by lipophagy, a specialised lysosomal degradative pathway of autophagy (Liu and Czaja, 2013, Martinez-Lopez and Singh, 2015). LD-localised ATGL contains LC3-interacting region motifs and promote lipophagy by associating to the autophagosome (Martinez-Lopez et al., 2016). Thus, we analysed the expression levels of *atgl-1* mRNA by quantitative PCR. We found that *atgl-1* expression increases in WT but not in *xbp-1(lof)* animals upon UPR^LP^ (Fig. 5F). These observations suggest that ATGL-1 may, in part, mediate lipid droplet clearance during UPR^LP^ and that functional IRE-1/XBP-1 axis is required in order to clear lipid droplets during lipid perturbation.

We next separated neutral lipids to investigate the abundance of TG and FFA in WT and UPR mutant animals subjected to *pmt-2* RNAi. We were expecting a large increased in TG based on our observation of LD increasing during UPR^LP^ (Fig. 2C). However, no significant increase in TG was observed at the exception of *atf-6(lof)* upon *pmt-2* RNAi treatment (Fig. S7A, B). Similarly, we did not observed any significant differences in FFA levels during lipid perturbation in WT and UPR mutants (Fig. S7C). These inconclusive findings may be due to (1) insignificant LP-induced accumulation of FFAs in comparison to the total pool of FFAs, (2) rapid reincorporation of FFAs into new lipids and breakdown by β-oxidation, or (3) LD-induced accumulation of FFAs in a subset of *C. elegans* cells. How these pathways contribute to FFA remodelling will be the challenge of future studies.

## DISCUSSION

Lipid perturbation refers to excessive accumulation of lipids in tissues including liver, pancreas and adipose (Hotamisligil and Erbay, 2008, Rinella and Sanyal, 2015). Dysfunctional UPR and apoptotic pathways ultimately lead to a disease outcome from this lipotoxicity. To better understand the role of the UPR during lipid perturbation and the consequence of a partial UPR programme, several studies have been conducted, focusing on their interconnections. As it is required for normal fatty acid synthesis as well as the regulation to the assembly and secretion of very low-density lipoprotein (VLDL), *XBP1* ablation leads to hypolipidemia in mice due to an abnormal decrease in plasma TG and cholesterol (Lee et al., 2008, So et al., 2012, Wang et al., 2012). High dietary carbohydrate is sufficient to increase FAs and cholesterol synthesis through XBP1 (Lee et al., 2008). Consequently, XBP1 is required to channel excess carbohydrate into lipids as its absence leads to insulin resistance in obese mice (Ozcan et al., 2004). XBP1 is also required to modulate phospholipid synthesis in an attempt to expand the ER membrane during proteotoxic stress to accommodate the increased load of misfolded proteins (Sriburi et al., 2004). The UPR modulates lipid metabolism-related genes and the absence of its sole regulator in yeast, Ire1, confers auxotrophy of inositol, a building block of phospholipids (Cox et al., 1993). We have previously shown that Ire1 is essential for cell survival during lipid perturbation, thereby highlighting the important role of the UPR to overcome lipotoxicity in yeast (Thibault et al., 2012). The UPR sensor PERK also plays a role in pathogenicity from lipotoxicity. Lipotoxicity-induced CHOP, the downstream target gene of PERK, promotes hepatic inflammation by activating the NK-κB pathway, thus promoting NASH and type 2 diabetes (Cunha et al., 2008, Willy et al., 2015). The ablation of ATF6, the third UPR sensor, induces NASH due to dysregulated lipid biosynthesis in mice under UPR^PT^ from tunicamycin treatment (Yamamoto et al., 2010). Thus, the three branches of the UPR are intimately linked to lipid homeostasis but their respective roles upon lipid perturbation in comparison to proteotoxic stress is still elusive. Here, we took a systematic global approach to determine genes that are UPR-regulated specifically from lipid perturbation.

To introduce lipid perturbation in *C. elegans*, we opted to genetically attenuate *pmt-2* which is required for the *de novo* PC biosynthesis (Fig. 1A). A similar approach has been used by other groups to mimic the physiological conditions associated to NAFLD in *C. elegans* (Walker et al., 2011, Ding et al., 2015, Smulan et al., 2016). Both required for *de novo* PC biosynthesis, *pmt-2* and *sams-1* depletion leads to lipid perturbation in worms (Li et al., 2004). Decrease of hepatic PC in mice (Walkey et al., 1998, Ozcan et al., 2004, Li et al., 2006, Fu et al., 2011a) and dietary deficiency of choline in humans are both associated to hepatic steatosis (Buchman et al., 1995, Gao et al., 2016). Initially, we subjected young adult worms to *pmt-2* RNAi for two days. However, no significant decrease in PC level was observed (data not shown). This could be due to the slow turnover of phospholipids in *C. elegans*. The absence of cell division in adult worms might not require the rapid synthesis of new membrane lipids thus genetic ablation of *pmt-2* may have little or no effect on PC levels (Kipreos, 2005). Thus, L1 stage worms treated with *pmt-2* RNAi were utilised as our UPR^LP^ model instead. Nonetheless, these conditions were sufficient to drastically induce lipid perturbation, lipid storage, and to strongly activate the UPR, all hallmark of NAFLD (Fig. 1, 2, S1-S3). Using this approach, we interrogated the role of each UPR branch during lipid perturbation-induced ER stress.

We employed *C. elegans* for its well-conserved UPR pathways and relative simplicity for genetic analysis. We examined the individual effects of *atf-6*, *ire-1*, and *pek-1* deficiency *in vivo*. Interestingly, ER stress induced by unfolded protein accumulation and lipid perturbation is found to be distinct from each other (Hou et al., 2014, Lajoie et al., 2012). The global transcriptomic analysis of UPR mutants subjected to lipid perturbation in comparison to proteotoxic-induced ER stress in WT allowed us to identify genes that are specifically regulated from the UPR^LP^ but not the UPR^PT^ (Fig. 3). To our knowledge, this is the first report identifying specific UPR-regulated genes induced specifically from lipid perturbation and not from proteotoxic stress. Our data show that a number of genes regulated by the UPR transducers are specific to lipid perturbation while a smaller number of genes are commonly modulated in proteotoxic and lipid stress. As expected, lipid perturbation-induced ER stress leads to altered regulation of transcription and protein modification processes through ATF-6, IRE-1, and PEK-1 (Fig. 3, 4). IRE-1 is the most conserved UPR transducer from yeast to mammals and it regulates the largest number of genes among the three UPR transducers (Fig. 3B). Consequently, we observed a severe phenotype in *ire-1;pmt-2(RNAi)* in comparison to *atf-6;pmt-2(RNAi)* and *pek-1;pmt-2(RNAi)* animals which both appear to be similar to WT animals.

Our autophagy screening revealed that autophagy is essential during lipid perturbation, suggesting its important role in regulating lipid metabolism. The change in lipid landscapes is at least partially mediated by autophagy, through the IRE-1/XBP-1 axis (Fig. 4, 5). Generally considered a cytoprotective response, autophagy can be modulated from ER stress. PERK has been reported to modulate autophagy by phosphorylating eIF2α resulting in a general translational inhibition (Matsumoto et al., 2013, Avivar-Valderas et al., 2011, Fujita et al., 2007, Kouroku et al., 2007). In parallel, PERK was also reported to regulate autophagy through the transcription factor ATF4 (orthologue of ATF-5) (Carra et al., 2009, Dever, 2002, Talloczy et al., 2002). Likewise, IRE1 modulates autophagy, independently of XBP1, by activating the Jun N-terminal kinase (JNK) pathway (Younce and Kolattukudy, 2012, Vidal et al., 2012, Pattingre et al., 2009, Wei et al., 2008b, Wei et al., 2008a, Ogata et al., 2006). Autophagy has also been reported to be activated (Younce and Kolattukudy, 2012) or inhibited (Vidal et al., 2012) from the IRE1/XBP1 axis (Adolph et al., 2013, Zhao et al., 2013, Hetz et al., 2009). Here, we have identified that autophagy is specifically upregulated from the IRE-1/XBP-1 axis upon UPR^LP^. We propose a model in which a reduction of PC activates the UPR which is essential to avert lipotoxicity (Fig. 6). This phenomenon is likely mediated by autophagy, where the process of recycling enlarged lipid droplets liberates FFAs stored within the droplets. This process of breaking down TG into FFAs might be required to rapidly repackage FFAs into deficient phospholipids in an attempt to reach lipid homeostasis at the ER. However as NAFLD can be caused from PC deficiency, this process will fail to restore lipid homeostasis leading to chronic or unresolved ER stress and may eventually lead to apoptosis, a characteristic of NAFLD and NASH (Walkey et al., 1998).

## MATERIALS AND METHODS

### Statistics

Error bars indicate standard error of the mean (SEM), calculated from at least three biological replicates, unless otherwise indicated. *P* values were calculated using one-way ANOVA with Tukey’s post hoc test, unless otherwise indicated and reported as *P* values with 4 significant digits in the figures. All statistical tests were performed using GraphPad Prism 7 software.

### *C. elegans* strains and RNAi constructs

All strains were grown at 20°C using standard *C. elegans* methods as previously described (Brenner, 1974, Stiernagle, 2006). Nematode growth media (NGM) agar plates were seeded with *Escherichia coli* strain OP50 for normal growth and with HT115 bacteria for RNAi feeding. RNAi feeding was performed as previously described (Timmons and Fire, 1998) and RNAi library was obtained from the Fire lab (Fire et al., 1998). The plasmids were sequenced to confirm their identity. Wild-type N2 Bristol, *atf-6*(ok551), *ire-1*(ok799), *pek-1*(ok275), *pmt-2*(vc1952), *hsp4::GFP*(sj4005) and *xbp-1(lof);hsp-4::GFP*(SJ17) were obtained from Caenorhabditis Genetic Center (CGC). *atf-6(lof)*;*hsp-4::GFP*, *ire-1(lof)*;*hsp-4::GFP* and *pek-1(lof)*;*hsp-4::GFP* were obtained by crossing *atf-6*(ok551), *ire-1*(ok799) and *pek-1*(ok275) to *hsp-4::GFP*(sj4005) as previously described (Fay, 2006).

### RNAi by feeding

RNAi was carried out as previously described (Timmons and Fire, 1998). Briefly, HT115 bacteria harbouring pL4440 plasmids were grown in LB medium containing 100 µg/ml ampicillin at 37°C until log phase (OD_600_ 0.6) and seeded onto NGM agar plates containing 50 µg/ml carbenicillin and 1 mM isopropyl β-D-1-thiogalactopyranoside (IPTG). Gravid adult worms were treated with hypochlorite and eggs were allowed to hatch overnight in M9 medium at 20°C to obtain L1 synchronised worms. Hatched L1 larvae were transferred to RNAi agar plates and grown until L4 larval to young adult stages. L4/young adult worms were harvested and incubated with 25 µg/ml tunicamycin in M9 medium for 4 hours at 20°C followed by M9 washes when indicated. To measure body length, worms were transferred to a 6 cm NGM plates without bacteria. Bright field images were acquired with a dissecting microscope (Nikon SMZ1500) fitted with a JVC digital camera at 100X magnification. Length measurements were performed in Fiji with WormSizer plugin (Moore et al., 2013).

### Quantitative real-time PCR

Ten thousand L4/young adult worms were collected, resuspended in water and lysed with a motorised pestle homogeniser. Total RNA was isolated using TRIzol reagent (Thermo Fisher, Waltham, MA) and subsequently purified using RNeasy Mini (Qiagen, Venlo, Netherlands) columns following manufacturer’s protocols. DNase treatment on column was carried out with RNase-Free DNase (Qiagen, Venlo, Netherlands) following manufacturer’s protocol. cDNA was synthesised from 2 μg of total RNA using RevertAid Reverse Transcriptase (Thermo Fisher, Waltham, MA) following manufacturer’s protocol. SYBR Green quantitative real-time PCR (qPCR) experiments were performed following manufacturer’s protocol using a QuantStudio 6 Flex Real-time PCR system (Applied Biosystems, Waltham, MA). Thirty nanograms of cDNA and 50 nM of paired-primer mix were used for each reaction. Relative mRNA was determined with the comparative Ct method (ΔΔCt) normalised to housekeeping gene *act-1*. Primers for qPCR are available upon request.

### Lipid extraction and phospholipid analysis

Approximately 10,000 L4 to young adult worms were harvested and washed thoroughly with M9 buffer, lysed with 1 mm silica beads by bead beating and subsequently lyophilised overnight (Virtis). All subsequent steps were carried out at 4°C. Total lipids were extracted from dried samples with chloroform:methanol (2:1) and concentrated. Total lipid extracts and POPE (1-palmitoyl-2-oleoyl-sn-glycero-3-phosphoethanolamine; 16:0-18:1n9 PE; Avanti Polar Lipids, Alabaster, AL) / DOPC (1,2-dioleoyl-sn-glycero-3-phosphocholine; 18:1n9 PC; Avanti Polar Lipids, Alabaster, AL) standard mix were spotted on HPTLC Silica gel 60 plates using Linomat 5 (CAMAG) and separated with chloroform:methanol:acetic acid:acetone:water (35:25:4:14:2). Phospholipids were visualised under long-wave ultraviolet (λ = 340 nm) by spraying 0.05 mg/ml of primuline dye in acetone:water (80:20) to the dried plates. Spots corresponding to PE and PC were scraped off the silica plates and transferred into 2 ml glass tubes. One hundred microliters of 1 mM C15:0 (pentadecanoic acid) was added to the tubes containing silica-bound phospholipids as an internal standard. The phospholipids were hydrolysed and esterified to fatty acid methyl esters (FAME) with 300 μl of 1.25 M HCl-methanol for 1 h at 80°C. FAMEs were extracted three times with 1 ml of hexane. Combine extracts were dried under nitrogen, resuspended in 100 μl hexane. FAMEs were separated by gas chromatography with flame ionization detector (GC-FID) (GC-2014; Shimadzu, Kyoto, Japan) using an ULBON HR-SS-10 50 m x 0.25 mm column (Shinwa, Tokyo, Japan). Supelco 37 component FAME mix was used to identify corresponding fatty acids (Sigma-Aldrich, St. Louis, MO). Data were normalised using internal standard C15:0 and worm dry weight.

### Lipid extraction and neutral lipid analysis

Around 35,000 worms treated on *pmt-2* RNAi for 48 hours were washed off the plate, spun down at 500 x *g* for one minute, and washed three times with 1x PBS/0.001% Triton X-100 wash solution. The worms were resuspended in 400 µL wash solution and lysed in 1 mm silica beads by bead beating. A 50 µL worm lysate was aliquoted from the 400 µL lysate for protein quantification using the BCA method. Worm lysates equalled to 150 µg of total soluble proteins were used for neutral lipid extraction following published method (Kniazeva et al., 2003). Briefly, 1.4 mL of ice-cold chloroform:methanol:formic acid (10:10:1) was added to the worm lysates in 4 mL glass tubes and vortexed vigorously for 2 minutes. Then, 2.1 mL chloroform:methanol:water (10:10:2) and vortexed vigorously for 2 minutes again. The tubes were stored at −20°C overnight for extraction. The tubes were spun at 3000 x *g* for 5 minutes for phase separation and the total lipid containing lower chloroform layer was extracted to new tubes and dried under nitrogen flow at room temperature. Dried total lipids were resuspended in 50 µL dichloromethane and spotted onto a silica plate as described above. TAG and FFA (Sigma) standards were loaded alongside the TLC plates. Neutral lipids were separated in a TLC tank with solvent consisted of hexane:diethyl ether:acetic acid at 75 mL:25 mL: 2 mL ratio. To visualize the lipids on the TLC plate, dried TLC plate was sprayed with 10% cupric sulphate (Sigma) in 8% phosphoric acid (Merck) and dried before development of the bands at a 120°C oven for 20 mins. The TLC plate was then scanned (Epson) and the bands intensity were quantified with ImageJ (NIH).

### Lipid droplets analysis by Sudan Black

Following RNAi feeding, fat storage of L4 worms was stained with Sudan Black B (Sigma-Aldrich, St. Louis, MO) as described previously with few modifications (Ogg and Ruvkun, 1998). Briefly, worms were fixed in 1% paraformaldehyde in M9 buffer for 30 minutes at room temperature, followed by three freeze-thaw cycles using liquid nitrogen. Worms were washed once with M9 and gradually dehydrated with 25%, 50% and 70% ethanol. Subsequently, fixed worms were stained with 50% saturated Sudan Black B in 70% ethanol (filtered with 0.22 µm membrane) for 30 minutes at room temperature with rocking. Stained worms were washed once with 25% ethanol for 30 minutes with rocking. Worms were mounted on a 2% agarose pads for imaging. Brightfield images of worms were taken with DMi8 inverted epifluorescence microscope (Leica, Wetzlar, Germany) with 20x and 63x objective lenses. To quantify lipid droplets, TIFF images taken at 63x magnification were converted to 8-bit grayscale, followed by background subtraction and thresholding was set using Fiji imaging software. The *Analyse particles* function was used to acquire mean corresponding to Sudan Black B intensity per worm.

### Fluorescence microscopy

To quantify GFP expression, transgenic line expressing GFP driven from *hsp-4* promoter was used. Worms were immobilised with 25 mM tetramisole and mounted on 2% agarose pad. Images were captured using DMi8 inverted epifluorescence microscope (Leica, Wetzlar, Germany) with 10x HC PL objective lens. Samples were excited at 488 nm and emission from 537 to 563 nm was collected. Three micrometers Z-stacks sections were merged and total fluorescence were quantified using Fiji imaging software.

### DNA microarray

Three independent populations of WT, *atf-6(lof), ire-1(lof), pek-1(lof)* and *pmt-2(lof)* worms were synchronised by hypochlorite treatment. L1 stage animals were treated with *pmt-2* RNAi or pL4440 empty vector for 48 h. Next, total RNA from the treated worms was isolated as described above. RNA quality control for microarray analysis was carried out using Agilent 2100 Bioanalyzer (Agilent Technologies, Santa Clara, CA). RNA samples with RIN score > 9.5 were deemed suitable for microarrays. The cDNAs were then synthesised from 100 ng of total RNA, purified, fragmented and hybridised to GeneChip C. elegans Gene 1.0 ST Arrays. Differentially expressed genes were identified using Affymetrix Transcriptome Analysis Console (TAC) 3.0 software. Threshold for selecting differentially expressed genes were set at a difference of more than 1.5-fold and one-way between subjects ANOVA *P* value < 0.05. GOrilla (http://cbl-gorilla.cs.technion.ac.il/) (Eden et al., 2009), REViGO (http://revigo.irb.hr/) (Supek et al., 2011), and DAVID (https://david.ncifcrf.gov/) (Huang et al., 2007) were used for GO terms analysis. Heat maps in the figures were generated using R Studio. Venn diagrams were generated using the following generator (http://bioinfo.genotoul.fr/jvenn/example.html).

For gene expression analysis, normalised and log transformed array data were imported to Cluster 3.0 for fold cut-off and hierarchical clustering. Genes were filtered to obtain those with a significant change in gene expression (fold change > 1.5 between RNAi treated and untreated samples and *P* < 0.05). The filtered data set was hierarchically clustered based on average linkage and Pearson correlation method, and the output was displayed in TreeView.

### RNAi screening of autophagy-related genes

RNAi screen was carried out as previously described with minor modifications (Lehner et al., 2006). Briefly, L1 larval stage animals were synchronised by hypochlorite treatment and exposed to *pmt-2 RNAi* on NGM agar plates for 36 h. Thereafter, worms were washed thrice with M9 and five to ten worms were seeded into a 96-well plate containing RNAi clones of autophagy-related functions. Control RNAi plates comprised worms exposed to pL4440 empty vector for 48 h and subsequently seeded into 96-well plates containing the same RNAi clones as above. Phenotypes of the worms were monitored over a five-day period. Phenotypes were compared to control RNAi plates where the worms were scored with sterility and reduced body size semi-quantitatively on a scale from 0 (wild-type) to 2 (100% sterility or stunted growth) (Lehner et al., 2006).

## SUPPLEMENTAL INFORMATION

Supplemental information includes seven figures, five Supplementary files and Supplemental References.

## AUTHOR CONTRIBUTIONS

Conceptualisation: G.T; Methodology: K.J.H., W.L., C.B.C., and G.T.; Formal Analysis: K.J.H.; Investigation: K.J.H., W.L., and C.B.C.; Resources: K.J.H. and W.L.; Writing – Original Draft: K.J.H. and G.T.; Writing – Review & Editing: K.J.H., W.L., C.B.C., and G.T.; Funding Acquisition: K.J.H. and G.T.

## ACKNOWLEDGEMENTS

We are grateful to Drs Fumio Motegi, Jean-Claude Labbé and Ronen Zaidel-Bar for providing reagents and technical support to introduce *C. elegans* as a new model system in the laboratory. Some strains were provided by the CGC, which is funded by NIH Office of Research Infrastructure Programs (P40 OD010440). We thank Peter Shyu Jr. for critical reading of the manuscript. This work was supported by the Nanyang Assistant Professorship programme from Nanyang Technological University and the Nanyang Technological University Research Scholarship to J.H.K. (predoctoral fellowship).

## SUPPLEMENTAL INFORMATION

### INVENTORY OF SUPPLEMENTAL INFORMATION

#### SUPPLEMENTAL FIGURES

**Figure S1.**
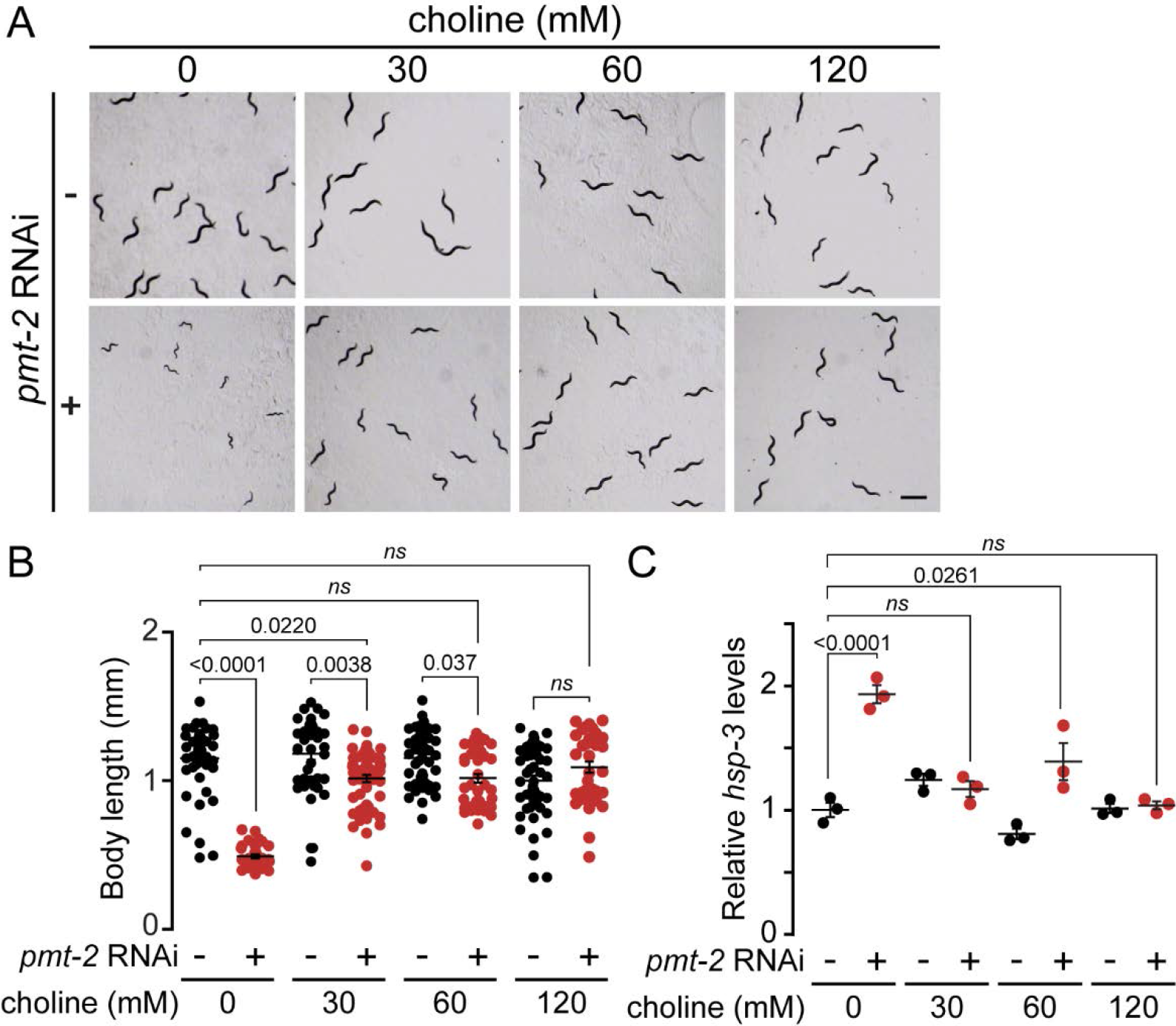
Refers to Figure 1. Choline supplementation restores developmental defects in *pmt-2(RNAi)* animals. (A) Representative choline rescue assay of *pmt-2(RNAi)* development defect. L1 worms were grown on *pmt-2* RNAi or empty vector on plate supplemented with 0, 30, 60, or 120 mM choline and images were taken after 48 h. Scale bar, 100 µm. (B) Body length quantification of worms from **A**. n > 24; ns, non-significant. (C) qPCR of *hsp-3* expression level in WT worms treated as in **A**. *ns* = non-significant.

**Figure S2.**
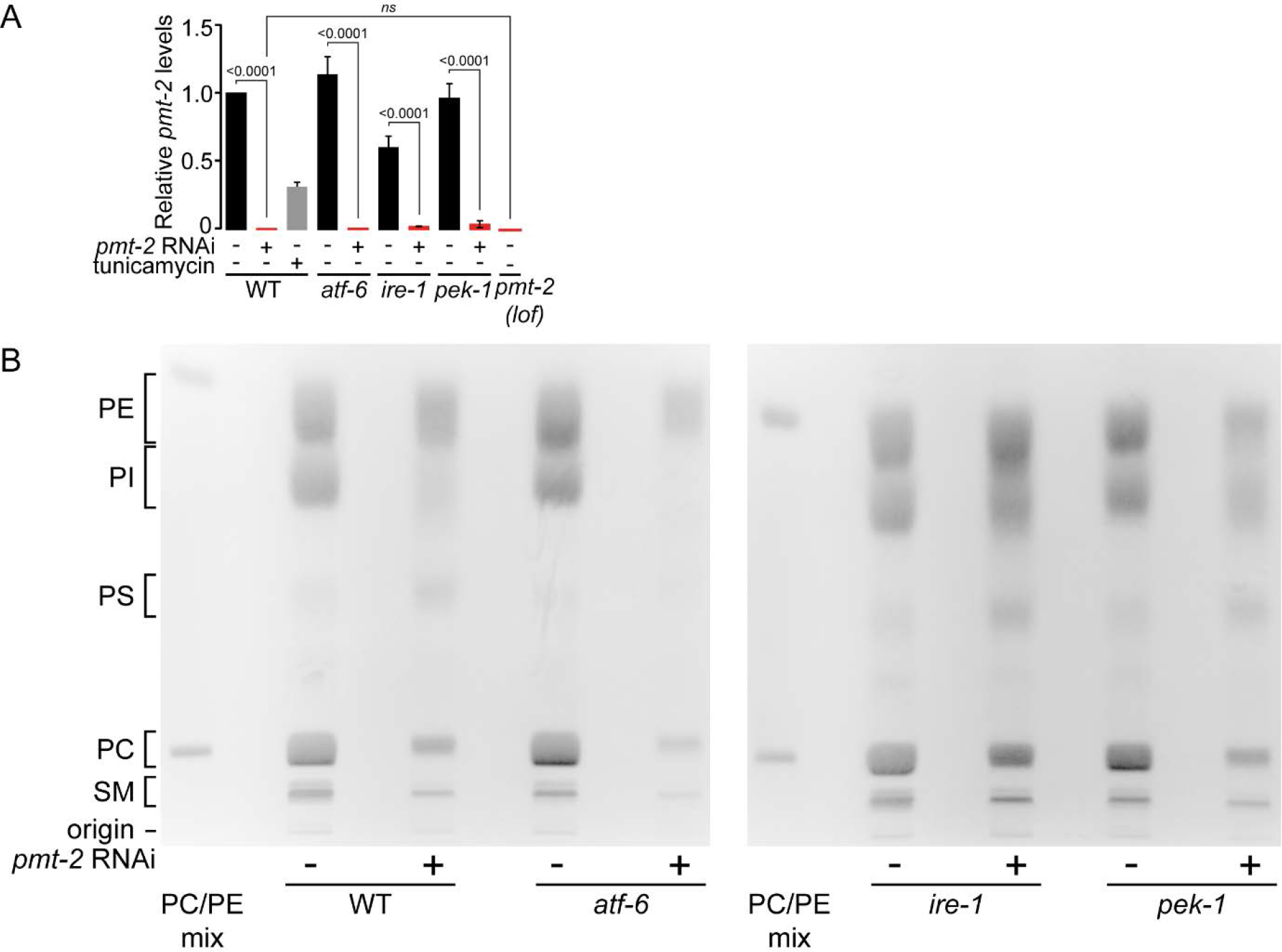
Refers to Figure 2A-B. Inactivation of *pmt-2* decreased PC content in worms. (A) qPCR of *pmt-2* expression after *pmt-2* RNAi treatment in WT, *atf-6(lof)*, *ire-1(lof),* and *pek-1(lof)* worms. *pmt-2(lof)* worms were used as a control. *pmt-2* RNAi treatment efficiently silenced expression of *pmt-2* across all the strains tested. *ns* = non-significant. (B) Representative separation of PE, MMPE, DMPE, and PC from total lipid extract using thin-layer chromatography (TLC). Comparison of phospholipid levels in WT, *atf-6(lof)*, *ire-1(lof)* and *pek-1(lof)* animals treated with *pmt-2* RNAi. POPE (1-palmitoyl-2-oleoyl-sn-glycero-3-phosphoethanolamine; 16:0-18:1n9 PE) and DOPC (1,2-dioleoyl-sn-glycero-3-phosphocholine; 18:1n9 PC) were used as markers.

**Figure S3.**
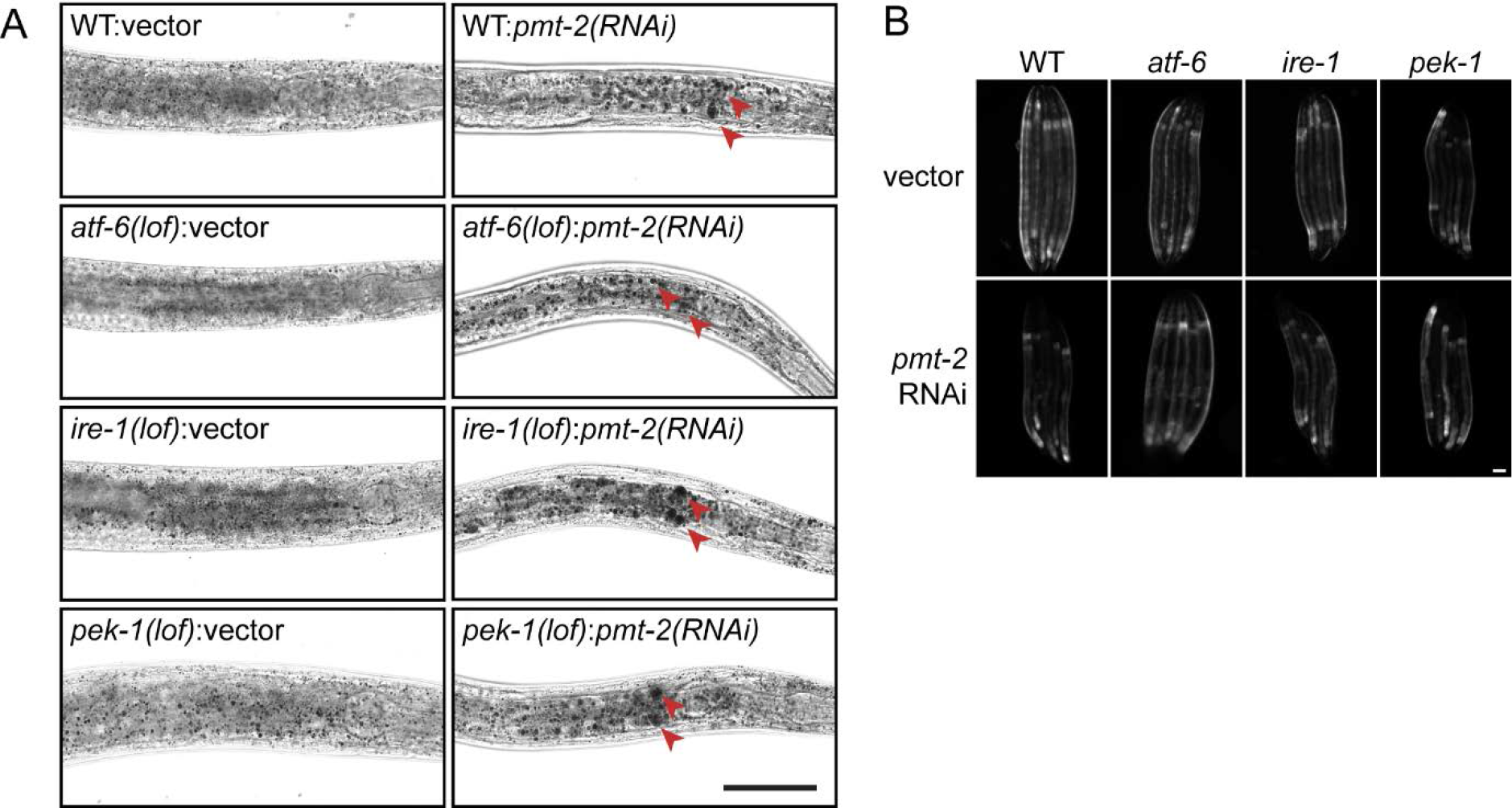
Refers to Figure 2C. Lipid perturbation induces lipid droplets accumulation and the UPR. (A) Representative images of lipid droplet visualised using Sudan Black B staining of WT, *atf-6(lof)*, *ire-1(lof)* and *pek-1(lof)* animals treated with *pmt-2* RNAi. Brightfield images of stained worms are shown using 63X objective lens. Scale bar, 100 µm. (B) Representative fluorescence images of *hsp-4::GFP* WT, *atf-6(lof)*, *ire-1(lof)* and *pek-1(lof)* animals treated with *pmt-2* RNAi. Scale bar, 100 µm.

**Figure S4.**
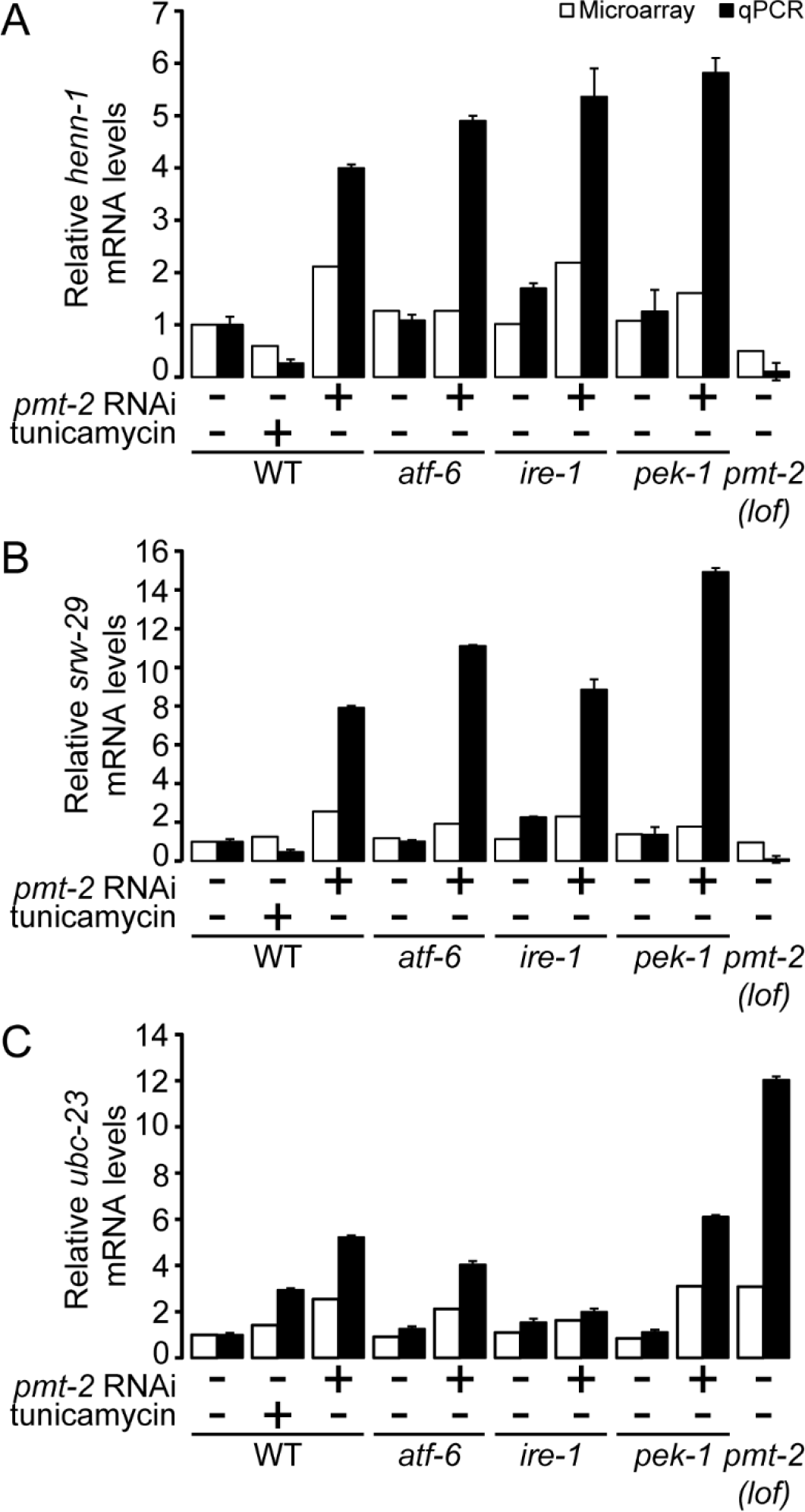
Refers to Figure 3. Validation of DNA microarray analysis using quantitative real-time PCR. (A-C) Comparison of *henn-1* (A), *srw-29* (B), and *ubc-23* (C) gene expression in WT, *atf-6(lof)*, *ire-1(lof)* and *pek-1(lof)* animals treated with *pmt-2* RNAi by DNA microarray (white bars) and qPCR (black bars).

**Figure S5.**
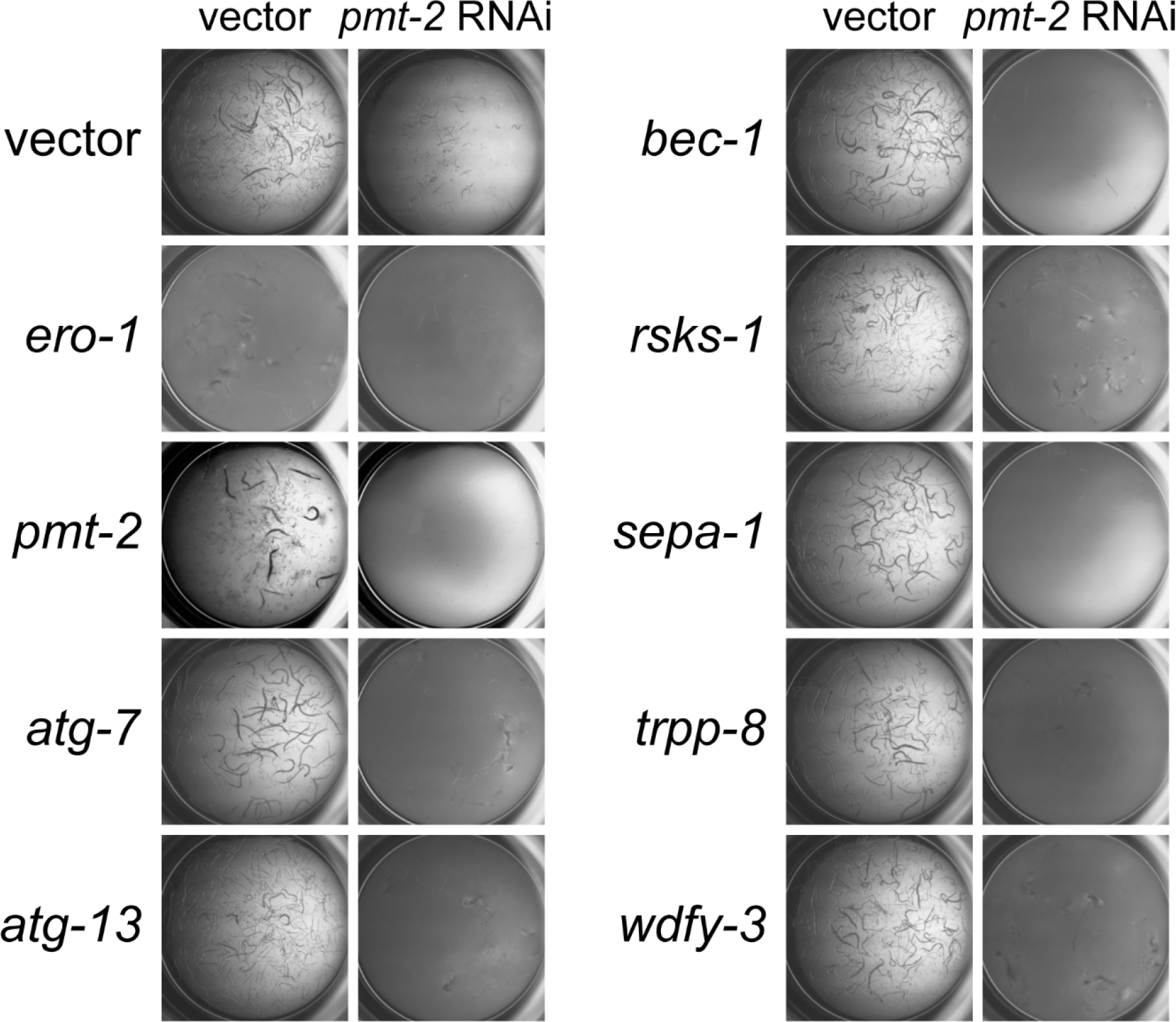
Refers to Figure 5. Autophagy is essential during lipid perturbation. WT worms were treated with vector or *pmt-2* RNAi for 36 h and subsequently subjected to RNAi in liquid media in a 96-well plate for 5 days. The worms were scored based on their developmental defects as described in Fig. 5A. *ero-1* RNAi was included as a positive control.

**Figure S6.**
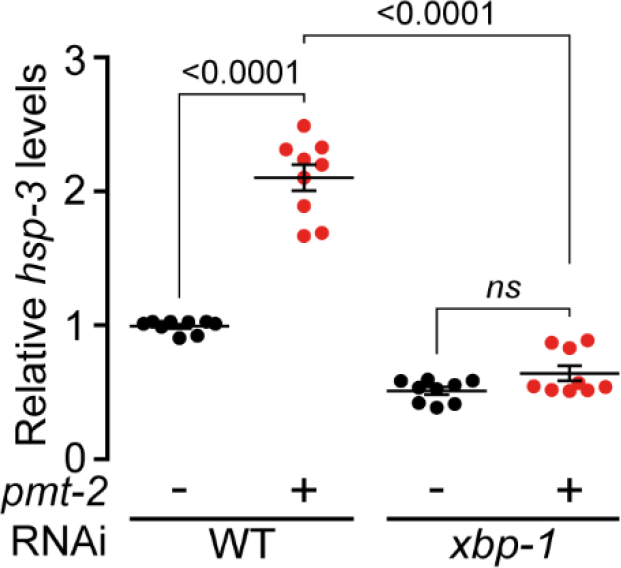
Refers to Figure 5. *hsp-3* is activated in an *xbp-1*-depedent manner in *pmt-2(RNAi)* worms. qPCR comparing expression of *hsp-3* in WT and *xbp-1(lof)* mutants treated with *pmt-2* RNAi. ns, non-significant.

**Figure S7.**
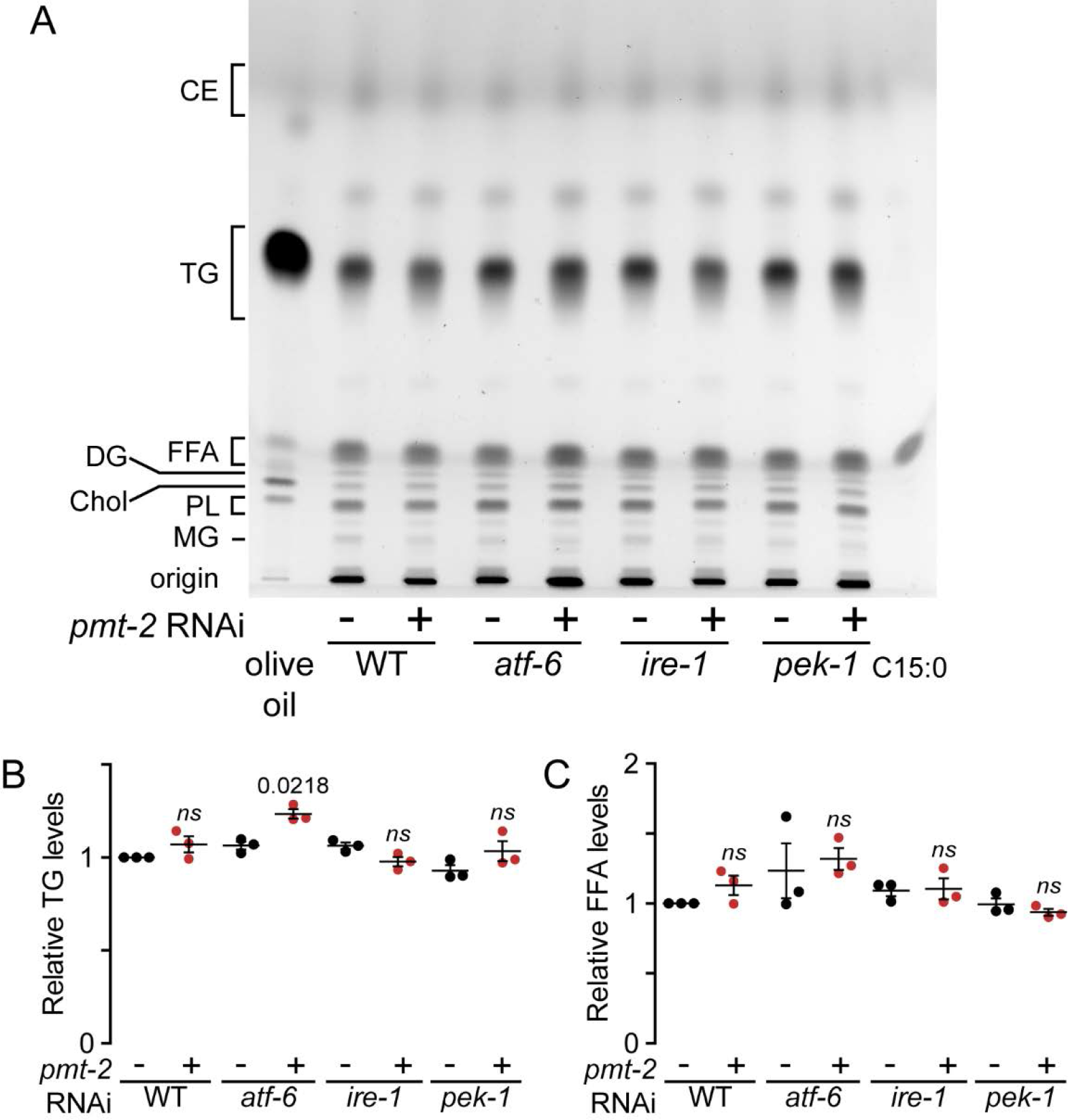
Refers to Figure 5. Neutral lipid separation in *pmt-2(RNAi)* worms. (A) Representative separation of neutral lipids from WT, *atf-6(lof)*, *ire-1(lof)* and *pek-1(lof)* animals treated with *pmt-2* RNAi. CE, cholesterol ester; DG, diacylglycerol; FFA, free fatty acid; MG, monoacylglycerol; PL, phospholipid; TG, triacylglycerol; olive oil and C15:0 pentadecanoic acid were added as markers of TG and FFA, respectively. Comparison of TG (B) and FFA (C) levels in WT, *atf-6(lof)*, *ire-1(lof)* and *pek-1(lof)* animals treated with *pmt-2* RNAi separated by TLC. FFA, free fatty acid; TG, triacylglycerol. One way ANOVA compared *pmt-2* RNAi treated to corresponding untreated mutant worm. ns, non-significant

**Supplementary file 1**, Refers to Figure 3A. List of upregulated and downregulated genes in *pmt-2(RNAi)* and WT treated with tunicamycin compared to WT animals. Excel Spreadsheet

**Supplementary file 2**, Refers to Figure 3B. **List of upregulated genes from the four-way Venn diagram.** Excel Spreadsheet

**Supplementary file 3**, Refers to Figure 3C. **Hierarchical clustering gene list.** Excel Spreadsheet

**Supplementary file 4**, Refers to Figure 4. **Predominant GO terms of each cluster.** Excel Spreadsheet

**Supplementary file 5**, Refers to Figure 5. **Phenotype from RNAi screen of autophagy genes**. Excel Spreadsheet

## REFERENCES

1. Adolph, T. E., Tomczak, M. F., Niederreiter, L., Ko, H. J., Bock, J., Martinez-Naves, E., Glickman, J. N., Tschurtschenthaler, M., Hartwig, J., Hosomi, S., Flak, M. B., Cusick, J. L., Kohno, K., Iwawaki, T., Billmann-Born, S., Raine, T., Bharti, R., Lucius, R., Kweon, M. N., Marciniak, S. J., Choi, A., Hagen, S. J., Schreiber, S., Rosenstiel, P., Kaser, A. & Blumberg, R. S. 2013. Paneth cells as a site of origin for intestinal inflammation. Nature, 503, 272–6.

2. Agouni, A., Mody, N., Owen, C., Czopek, A., Zimmer, D., Bentires-Alj, M., Bence, K. K. & Delibegovic, M. 2011. Liver-specific deletion of protein tyrosine phosphatase (PTP) 1B improves obesity- and pharmacologically induced endoplasmic reticulum stress. Biochem J, 438, 369–78.

3. Arendt, B. M., Ma, D. W., Simons, B., Noureldin, S. A., Therapondos, G., Guindi, M., Sherman, M. & Allard, J. P. 2013. Nonalcoholic fatty liver disease is associated with lower hepatic and erythrocyte ratios of phosphatidylcholine to phosphatidylethanolamine. Appl Physiol Nutr Metab, 38, 334–40.

4. Avivar-Valderas, A., Salas, E., Bobrovnikova-Marjon, E., Diehl, J. A., Nagi, C., Debnath, J. & Aguirre-Ghiso, J. A. 2011. PERK integrates autophagy and oxidative stress responses to promote survival during extracellular matrix detachment. Mol Cell Biol, 31, 3616–29.

5. Bettaieb, A., Matsuo, K., Matsuo, I., Wang, S., Melhem, R., Koromilas, A. E. & Haj, F. G. 2012. Protein tyrosine phosphatase 1B deficiency potentiates PERK/eIF2alpha signaling in brown adipocytes. PLoS One, 7, e34412.

6. Braakman, I. & Bulleid, N. J. 2011. Protein folding and modification in the mammalian endoplasmic reticulum. Annu Rev Biochem, 80, 71–99.

7. Brendza, K. M., Haakenson, W., Cahoon, R. E., Hicks, L. M., Palavalli, L. H., Chiapelli, B. J., McLaird, M., McCarter, J. P., Williams, D. J., Hresko, M. C. & Jez, J. M. 2007. Phosphoethanolamine N-methyltransferase (PMT-1) catalyses the first reaction of a new pathway for phosphocholine biosynthesis in Caenorhabditis elegans. Biochem J, 404, 439–48.

8. Brenner, S. 1974. The genetics of Caenorhabditis elegans. Genetics, 77, 71–94.

9. Buchman, A. L., Dubin, M. D., Moukarzel, A. A., Jenden, D. J., Roch, M., Rice, K. M., Gornbein, J. & Ament, M. E. 1995. Choline deficiency: a cause of hepatic steatosis during parenteral nutrition that can be reversed with intravenous choline supplementation. Hepatology, 22, 1399–403.

10. Calfon, M., Zeng, H., Urano, F., Till, J. H., Hubbard, S. R., Harding, H. P., Clark, S. G. & Ron, D. 2002. IRE1 couples endoplasmic reticulum load to secretory capacity by processing the XBP-1 mRNA. Nature, 415, 92–6.

11. Carra, S., Brunsting, J. F., Lambert, H., Landry, J. & Kampinga, H. H. 2009. HspB8 participates in protein quality control by a non-chaperone-like mechanism that requires eIF2{alpha} phosphorylation. J Biol Chem, 284, 5523–32.

12. Cox, J. S., Shamu, C. E. & Walter, P. 1993. Transcriptional induction of genes encoding endoplasmic reticulum resident proteins requires a transmembrane protein kinase. Cell, 73, 1197–206.

13. Cox, J. S. & Walter, P. 1996. A novel mechanism for regulating activity of a transcription factor that controls the unfolded protein response. Cell, 87, 391–404.

14. Cunha, D. A., Hekerman, P., Ladriere, L., Bazarra-Castro, A., Ortis, F., Wakeham, M. C., Moore, F., Rasschaert, J., Cardozo, A. K., Bellomo, E., Overbergh, L., Mathieu, C., Lupi, R., Hai, T., Herchuelz, A., Marchetti, P., Rutter, G. A., Eizirik, D. L. & Cnop, M. 2008. Initiation and execution of lipotoxic ER stress in pancreatic beta-cells. J Cell Sci, 121, 2308–18.

15. Dever, T. E. 2002. Gene-specific regulation by general translation factors. Cell, 108, 545–56.

16. Ding, W., Smulan, L. J., Hou, N. S., Taubert, S., Watts, J. L. & Walker, A. K. 2015. s-Adenosylmethionine Levels Govern Innate Immunity through Distinct Methylation-Dependent Pathways. Cell Metab, 22, 633–45.

17. Doycheva, I., Watt, K. D., Rifai, G., Abou Mrad, R., Lopez, R., Zein, N. N., Carey, W. D. & Alkhouri, N. 2017. Increasing Burden of Chronic Liver Disease Among Adolescents and Young Adults in the USA: A Silent Epidemic. Dig Dis Sci, 62, 1373–1380.

18. Eden, E., Navon, R., Steinfeld, I., Lipson, D. & Yakhini, Z. 2009. GOrilla: a tool for discovery and visualization of enriched GO terms in ranked gene lists. BMC Bioinformatics, 10, 48.

19. Ericson, M. C., Gafford, J. T. & Elbein, A. D. 1977. Tunicamycin inhibits GlcNAc-lipid formation in plants. J Biol Chem, 252, 7431–3.

20. Fay, D. 2006. Genetic mapping and manipulation: chapter 1--Introduction and basics. WormBook, 1–12.

21. Fire, A., Xu, S., Montgomery, M. K., Kostas, S. A., Driver, S. E. & Mello, C. C. 1998. Potent and specific genetic interference by double-stranded RNA in Caenorhabditis elegans. Nature, 391, 806–11.

22. Fu, S., Yang, L., Li, P., Hofmann, O., Dicker, L., Hide, W., Lin, X., Watkins, S. M., Ivanov, A. R. & Hotamisligil, G. S. 2011a. Aberrant lipid metabolism disrupts calcium homeostasis causing liver endoplasmic reticulum stress in obesity. Nature, 473, 528–531.

23. Fu, S., Yang, L., Li, P., Hofmann, O., Dicker, L., Hide, W., Lin, X., Watkins, S. M., Ivanov, A. R. & Hotamisligil, G. S. 2011b. Aberrant lipid metabolism disrupts calcium homeostasis causing liver endoplasmic reticulum stress in obesity. Nature, 473, 528–31.

24. Fujita, E., Kouroku, Y., Isoai, A., Kumagai, H., Misutani, A., Matsuda, C., Hayashi, Y. K. & Momoi, T. 2007. Two endoplasmic reticulum-associated degradation (ERAD) systems for the novel variant of the mutant dysferlin: ubiquitin/proteasome ERAD(I) and autophagy/lysosome ERAD(II). Hum Mol Genet, 16, 618–29.

25. Gao, X., Wang, Y., Randell, E., Pedram, P., Yi, Y., Gulliver, W. & Sun, G. 2016. Higher Dietary Choline and Betaine Intakes Are Associated with Better Body Composition in the Adult Population of Newfoundland, Canada. PLoS One, 11, e0155403.

26. Gross, D. A. & Silver, D. L. 2014. Cytosolic lipid droplets: from mechanisms of fat storage to disease. Crit Rev Biochem Mol Biol, 49, 304–26.

27. Guo, Y., Walther, T. C., Rao, M., Stuurman, N., Goshima, G., Terayama, K., Wong, J. S., Vale, R. D., Walter, P. & Farese, R. V. 2008. Functional genomic screen reveals genes involved in lipid-droplet formation and utilization. Nature, 453, 657–61.

28. Han, J. & Kaufman, R. J. 2016. The role of ER stress in lipid metabolism and lipotoxicity. J Lipid Res, 57, 1329–38.

29. Harding, H. P., Zhang, Y., Zeng, H., Novoa, I., Lu, P. D., Calfon, M., Sadri, N., Yun, C., Popko, B., Paules, R., Stojdl, D. F., Bell, J. C., Hettmann, T., Leiden, J. M. & Ron, D. 2003. An integrated stress response regulates amino acid metabolism and resistance to oxidative stress. Mol Cell, 11, 619–33.

30. Hashemi, H. F. & Goodman, J. M. 2015. The life cycle of lipid droplets. Curr Opin Cell Biol, 33, 119–24.

31. Hetz, C., Thielen, P., Matus, S., Nassif, M., Court, F., Kiffin, R., Martinez, G., Cuervo, A. M., Brown, R. H. & Glimcher, L. H. 2009. XBP-1 deficiency in the nervous system protects against amyotrophic lateral sclerosis by increasing autophagy. Genes Dev, 23, 2294–306.

32. Ho, N., Xu, C. & Thibault, G. 2018. From the unfolded protein response to metabolic diseases - lipids under the spotlight. J Cell Sci, 131.

33. Hollien, J., Lin, J. H., Li, H., Stevens, N., Walter, P. & Weissman, J. S. 2009. Regulated Ire1-dependent decay of messenger RNAs in mammalian cells. J Cell Biol, 186, 323–31.

34. Horl, G., Wagner, A., Cole, L. K., Malli, R., Reicher, H., Kotzbeck, P., Kofeler, H., Hofler, G., Frank, S., Bogner-Strauss, J. G., Sattler, W., Vance, D. E. & Steyrer, E. 2011. Sequential synthesis and methylation of phosphatidylethanolamine promote lipid droplet biosynthesis and stability in tissue culture and in vivo. J Biol Chem, 286, 17338–50.

35. Hotamisligil, G. S. & Erbay, E. 2008. Nutrient sensing and inflammation in metabolic diseases. Nat Rev Immunol, 8, 923–34.

36. Hou, N. S., Gutschmidt, A., Choi, D. Y., Pather, K., Shi, X., Watts, J. L., Hoppe, T. & Taubert, S. 2014. Activation of the endoplasmic reticulum unfolded protein response by lipid disequilibrium without disturbed proteostasis in vivo. Proc Natl Acad Sci U S A, 111, E2271–80.

37. Huang, D. W., Sherman, B. T., Tan, Q., Collins, J. R., Alvord, W. G., Roayaei, J., Stephens, R., Baseler, M. W., Lane, H. C. & Lempicki, R. A. 2007. The DAVID Gene Functional Classification Tool: a novel biological module-centric algorithm to functionally analyze large gene lists. Genome Biol, 8, R183.

38. Jia, K., Hart, A. C. & Levine, B. 2007. Autophagy genes protect against disease caused by polyglutamine expansion proteins in Caenorhabditis elegans. Autophagy, 3, 21–5.

39. Kim, M. H., Aydemir, T. B., Kim, J. & Cousins, R. J. 2017. Hepatic ZIP14-mediated zinc transport is required for adaptation to endoplasmic reticulum stress. Proc Natl Acad Sci U S A, 114, E5805–E5814.

40. Kipreos, E. T. 2005. C. elegans cell cycles: invariance and stem cell divisions. Nat Rev Mol Cell Biol, 6, 766–76.

41. Kniazeva, M., Sieber, M., McCauley, S., Zhang, K., Watts, J. L. & Han, M. 2003. Suppression of the ELO-2 FA elongation activity results in alterations of the fatty acid composition and multiple physiological defects, including abnormal ultradian rhythms, in Caenorhabditis elegans. Genetics, 163, 159–69.

42. Kouroku, Y., Fujita, E., Tanida, I., Ueno, T., Isoai, A., Kumagai, H., Ogawa, S., Kaufman, R. J., Kominami, E. & Momoi, T. 2007. ER stress (PERK/eIF2alpha phosphorylation) mediates the polyglutamine-induced LC3 conversion, an essential step for autophagy formation. Cell Death Differ, 14, 230–9.

43. Lagace, T. A. & Ridgway, N. D. 2013. The role of phospholipids in the biological activity and structure of the endoplasmic reticulum. Biochim Biophys Acta, 1833, 2499–510.

44. Lajoie, P., Moir, R. D., Willis, I. M. & Snapp, E. L. 2012. Kar2p availability defines distinct forms of endoplasmic reticulum stress in living cells. Mol Biol Cell, 23, 955–64.

45. Lee, A. H., Scapa, E. F., Cohen, D. E. & Glimcher, L. H. 2008. Regulation of hepatic lipogenesis by the transcription factor XBP1. Science, 320, 1492–6.

46. Lehner, B., Tischler, J. & Fraser, A. G. 2006. RNAi screens in Caenorhabditis elegans in a 96-well liquid format and their application to the systematic identification of genetic interactions. Nat Protoc, 1, 1617–20.

47. Li, Y., Ge, M., Ciani, L., Kuriakose, G., Westover, E. J., Dura, M., Covey, D. F., Freed, J. H., Maxfield, F. R., Lytton, J. & Tabas, I. 2004. Enrichment of endoplasmic reticulum with cholesterol inhibits sarcoplasmic-endoplasmic reticulum calcium ATPase-2b activity in parallel with increased order of membrane lipids: implications for depletion of endoplasmic reticulum calcium stores and apoptosis in cholesterol-loaded macrophages. J Biol Chem, 279, 37030–9.

48. Li, Y., Na, K., Lee, H. J., Lee, E. Y. & Paik, Y. K. 2011. Contribution of sams-1 and pmt-1 to lipid homoeostasis in adult Caenorhabditis elegans. J Biochem, 149, 529–38.

49. Li, Z., Agellon, L. B., Allen, T. M., Umeda, M., Jewell, L., Mason, A. & Vance, D. E. 2006. The ratio of phosphatidylcholine to phosphatidylethanolamine influences membrane integrity and steatohepatitis. Cell Metab, 3, 321–31.

50. Li, Z., Agellon, L. B. & Vance, D. E. 2005. Phosphatidylcholine homeostasis and liver failure. J Biol Chem, 280, 37798–802.

51. Ling, J., Chaba, T., Zhu, L. F., Jacobs, R. L. & Vance, D. E. 2012. Hepatic ratio of phosphatidylcholine to phosphatidylethanolamine predicts survival after partial hepatectomy in mice. Hepatology, 55, 1094–102.

52. Ling, J., Reynolds, N. & Ibba, M. 2009. Aminoacyl-tRNA synthesis and translational quality control. Annu Rev Microbiol, 63, 61–78.

53. Liu, K. & Czaja, M. J. 2013. Regulation of lipid stores and metabolism by lipophagy. Cell Death Differ, 20, 3–11.

54. Long, X., Spycher, C., Han, Z. S., Rose, A. M., Muller, F. & Avruch, J. 2002. TOR deficiency in C. elegans causes developmental arrest and intestinal atrophy by inhibition of mRNA translation. Curr Biol, 12, 1448–61.

55. Martinez-Lopez, N., Garcia-Macia, M., Sahu, S., Athonvarangkul, D., Liebling, E., Merlo, P., Cecconi, F., Schwartz, G. J. & Singh, R. 2016. Autophagy in the CNS and Periphery Coordinate Lipophagy and Lipolysis in the Brown Adipose Tissue and Liver. Cell Metab, 23, 113–27.

56. Martinez-Lopez, N. & Singh, R. 2015. Autophagy and Lipid Droplets in the Liver. Annu Rev Nutr, 35, 215–37.

57. Matsumoto, H., Miyazaki, S., Matsuyama, S., Takeda, M., Kawano, M., Nakagawa, H., Nishimura, K. & Matsuo, S. 2013. Selection of autophagy or apoptosis in cells exposed to ER-stress depends on ATF4 expression pattern with or without CHOP expression. Biol Open, 2, 1084–90.

58. Melendez, A., Talloczy, Z., Seaman, M., Eskelinen, E. L., Hall, D. H. & Levine, B. 2003. Autophagy genes are essential for dauer development and life-span extension in C. elegans. Science, 301, 1387–91.

59. Mizushima, N. & Komatsu, M. 2011. Autophagy: renovation of cells and tissues. Cell, 147, 728–41.

60. Moore, B. T., Jordan, J. M. & Baugh, L. R. 2013. WormSizer: high-throughput analysis of nematode size and shape. PLoS One, 8, e57142.

61. Morein, S., Andersson, A., Rilfors, L. & Lindblom, G. 1996. Wild-type Escherichia coli cells regulate the membrane lipid composition in a “window” between gel and non-lamellar structures. J Biol Chem, 271, 6801–9.

62. Mota, M., Banini, B. A., Cazanave, S. C. & Sanyal, A. J. 2016. Molecular mechanisms of lipotoxicity and glucotoxicity in nonalcoholic fatty liver disease. Metabolism, 65, 1049–61.

63. Ng, B. S. H., Shyu, P. T., Chaw, R., Seah, Y. L. & Thibault, G. 2017. Lipid perturbation compromises UPR-mediated ER homeostasis as a result of premature degradation of membrane proteins. bioRxiv.

64. Ng, D. T., Spear, E. D. & Walter, P. 2000. The unfolded protein response regulates multiple aspects of secretory and membrane protein biogenesis and endoplasmic reticulum quality control. J Cell Biol, 150, 77–88.

65. Nguyen, T. B., Louie, S. M., Daniele, J. R., Tran, Q., Dillin, A., Zoncu, R., Nomura, D. K. & Olzmann, J. A. 2017. DGAT1-Dependent Lipid Droplet Biogenesis Protects Mitochondrial Function during Starvation-Induced Autophagy. Dev Cell, 42, 9–21 e5.

66. Novoa, I., Zeng, H., Harding, H. P. & Ron, D. 2001. Feedback inhibition of the unfolded protein response by GADD34-mediated dephosphorylation of eIF2alpha. J Cell Biol, 153, 1011–22.

67. Ogata, M., Hino, S., Saito, A., Morikawa, K., Kondo, S., Kanemoto, S., Murakami, T., Taniguchi, M., Tanii, I., Yoshinaga, K., Shiosaka, S., Hammarback, J. A., Urano, F. & Imaizumi, K. 2006. Autophagy is activated for cell survival after endoplasmic reticulum stress. Mol Cell Biol, 26, 9220–31.

68. Ogg, S. & Ruvkun, G. 1998. The C. elegans PTEN homolog, DAF-18, acts in the insulin receptor-like metabolic signaling pathway. Mol Cell, 2, 887–93.

69. Oursel, D., Loutelier-Bourhis, C., Orange, N., Chevalier, S., Norris, V. & Lange, C. M. 2007. Lipid composition of membranes of Escherichia coli by liquid chromatography/tandem mass spectrometry using negative electrospray ionization. Rapid Commun Mass Spectrom, 21, 1721–8.

70. Oyadomari, S., Koizumi, A., Takeda, K., Gotoh, T., Akira, S., Araki, E. & Mori, M. 2002. Targeted disruption of the Chop gene delays endoplasmic reticulum stress-mediated diabetes. J Clin Invest, 109, 525–32.

71. Ozcan, U., Cao, Q., Yilmaz, E., Lee, A. H., Iwakoshi, N. N., Ozdelen, E., Tuncman, G., Gorgun, C., Glimcher, L. H. & Hotamisligil, G. S. 2004. Endoplasmic reticulum stress links obesity, insulin action, and type 2 diabetes. Science, 306, 457–61.

72. Palavalli, L. H., Brendza, K. M., Haakenson, W., Cahoon, R. E., McLaird, M., Hicks, L. M., McCarter, J. P., Williams, D. J., Hresko, M. C. & Jez, J. M. 2006. Defining the role of phosphomethylethanolamine N-methyltransferase from Caenorhabditis elegans in phosphocholine biosynthesis by biochemical and kinetic analysis. Biochemistry, 45, 6056–65.

73. Pattingre, S., Bauvy, C., Carpentier, S., Levade, T., Levine, B. & Codogno, P. 2009. Role of JNK1-dependent Bcl-2 phosphorylation in ceramide-induced macroautophagy. J Biol Chem, 284, 2719–28.

74. Promlek, T., Ishiwata-Kimata, Y., Shido, M., Sakuramoto, M., Kohno, K. & Kimata, Y. 2011. Membrane aberrancy and unfolded proteins activate the endoplasmic reticulum stress sensor Ire1 in different ways. Mol Biol Cell, 22, 3520–32.

75. Puri, P., Baillie, R. A., Wiest, M. M., Mirshahi, F., Choudhury, J., Cheung, O., Sargeant, C., Contos, M. J. & Sanyal, A. J. 2007. A lipidomic analysis of nonalcoholic fatty liver disease. Hepatology, 46, 1081–90.

76. Rinella, M. E. & Sanyal, A. J. 2015. NAFLD in 2014: Genetics, diagnostics and therapeutic advances in NAFLD. Nat Rev Gastroenterol Hepatol, 12, 65–6.

77. Rubio, C., Pincus, D., Korennykh, A., Schuck, S., El-Samad, H. & Walter, P. 2011. Homeostatic adaptation to endoplasmic reticulum stress depends on Ire1 kinase activity. J Cell Biol, 193, 171–84.

78. Rutkowski, D. T., Arnold, S. M., Miller, C. N., Wu, J., Li, J., Gunnison, K. M., Mori, K., Sadighi Akha, A. A., Raden, D. & Kaufman, R. J. 2006. Adaptation to ER stress is mediated by differential stabilities of pro-survival and pro-apoptotic mRNAs and proteins. PLoS Biol, 4, e374.

79. Sathyanarayan, A., Mashek, M. T. & Mashek, D. G. 2017. ATGL Promotes Autophagy/Lipophagy via SIRT1 to Control Hepatic Lipid Droplet Catabolism. Cell Rep, 19, 1–9.

80. Schroder, M. & Kaufman, R. J. 2005. ER stress and the unfolded protein response. Mutat Res, 569, 29–63.

81. Shen, X., Ellis, R. E., Sakaki, K. & Kaufman, R. J. 2005. Genetic interactions due to constitutive and inducible gene regulation mediated by the unfolded protein response in C. elegans. PLoS Genet, 1, e37.

82. Shen, X., Zhang, K. & Kaufman, R. J. 2004. The unfolded protein response--a stress signaling pathway of the endoplasmic reticulum. J Chem Neuroanat, 28, 79–92.

83. Smulan, L. J., Ding, W., Freinkman, E., Gujja, S., Edwards, Y. J. K. & Walker, A. K. 2016. Cholesterol-Independent SREBP-1 Maturation Is Linked to ARF1 Inactivation. Cell Rep, 16, 9–18.

84. So, J. S., Hur, K. Y., Tarrio, M., Ruda, V., Frank-Kamenetsky, M., Fitzgerald, K., Koteliansky, V., Lichtman, A. H., Iwawaki, T., Glimcher, L. H. & Lee, A. H. 2012. Silencing of lipid metabolism genes through IRE1alpha-mediated mRNA decay lowers plasma lipids in mice. Cell Metab, 16, 487–99.

85. Sriburi, R., Jackowski, S., Mori, K. & Brewer, J. W. 2004. XBP1: a link between the unfolded protein response, lipid biosynthesis, and biogenesis of the endoplasmic reticulum. J Cell Biol, 167, 35–41.

86. Stiernagle, T. 2006. Maintenance of C. elegans. WormBook, 1-11.

87. Supek, F., Bosnjak, M., Skunca, N. & Smuc, T. 2011. REVIGO summarizes and visualizes long lists of gene ontology terms. PLoS One, 6, e21800.

88. Takacs-Vellai, K., Vellai, T., Puoti, A., Passannante, M., Wicky, C., Streit, A., Kovacs, A. L. & Muller, F. 2005. Inactivation of the autophagy gene bec-1 triggers apoptotic cell death in C. elegans. Curr Biol, 15, 1513–7.

89. Talloczy, Z., Jiang, W., Virgin, H. W. T., Leib, D. A., Scheuner, D., Kaufman, R. J., Eskelinen, E. L. & Levine, B. 2002. Regulation of starvation- and virus-induced autophagy by the eIF2alpha kinase signaling pathway. Proc Natl Acad Sci U S A, 99, 190–5.

90. Thibault, G., Ismail, N. & Ng, D. T. 2011. The unfolded protein response supports cellular robustness as a broad-spectrum compensatory pathway. Proc Natl Acad Sci U S A, 108, 20597–602.

91. Thibault, G., Shui, G., Kim, W., McAlister, G. C., Ismail, N., Gygi, S. P., Wenk, M. R. & Ng, D. T. 2012. The membrane stress response buffers lethal effects of lipid disequilibrium by reprogramming the protein homeostasis network. Mol Cell, 48, 16–27.

92. Timmons, L. & Fire, A. 1998. Specific interference by ingested dsRNA. Nature, 395, 854.

93. Tiniakos, D. G., Vos, M. B. & Brunt, E. M. 2010. Nonalcoholic fatty liver disease: pathology and pathogenesis. Annu Rev Pathol, 5, 145–71.

94. Travers, K. J., Patil, C. K., Wodicka, L., Lockhart, D. J., Weissman, J. S. & Walter, P. 2000. Functional and genomic analyses reveal an essential coordination between the unfolded protein response and ER-associated degradation. Cell, 101, 249–58.

95. Urano, F., Calfon, M., Yoneda, T., Yun, C., Kiraly, M., Clark, S. G. & Ron, D. 2002. A survival pathway for Caenorhabditis elegans with a blocked unfolded protein response. J Cell Biol, 158, 639–46.

96. Velazquez, A. P., Tatsuta, T., Ghillebert, R., Drescher, I. & Graef, M. 2016. Lipid droplet-mediated ER homeostasis regulates autophagy and cell survival during starvation. J Cell Biol, 212, 621–31.

97. Vidal, R. L., Figueroa, A., Court, F. A., Thielen, P., Molina, C., Wirth, C., Caballero, B., Kiffin, R., Segura-Aguilar, J., Cuervo, A. M., Glimcher, L. H. & Hetz, C. 2012. Targeting the UPR transcription factor XBP1 protects against Huntington’s disease through the regulation of FoxO1 and autophagy. Hum Mol Genet, 21, 2245–62.

98. Volmer, R. & Ron, D. 2015. Lipid-dependent regulation of the unfolded protein response. Curr Opin Cell Biol, 33, 67–73.

99. Volmer, R., Van der Ploeg, K. & Ron, D. 2013. Membrane lipid saturation activates endoplasmic reticulum unfolded protein response transducers through their transmembrane domains. Proc Natl Acad Sci U S A, 110, 4628–33.

100. Walker, A. K., Jacobs, R. L., Watts, J. L., Rottiers, V., Jiang, K., Finnegan, D. M., Shioda, T., Hansen, M., Yang, F., Niebergall, L. J., Vance, D. E., Tzoneva, M., Hart, A. C. & Naar, A. M. 2011. A conserved SREBP-1/phosphatidylcholine feedback circuit regulates lipogenesis in metazoans. Cell, 147, 840–52.

101. Walkey, C. J., Yu, L., Agellon, L. B. & Vance, D. E. 1998. Biochemical and evolutionary significance of phospholipid methylation. J Biol Chem, 273, 27043–6.

102. Walter, P. & Ron, D. 2011. The unfolded protein response: from stress pathway to homeostatic regulation. Science, 334, 1081–6.

103. Wang, C., Saar, V., Leung, K. L., Chen, L. & Wong, G. 2018. Human amyloid beta peptide and tau co-expression impairs behavior and causes specific gene expression changes in Caenorhabditis elegans. Neurobiol Dis, 109, 88–101.

104. Wang, S., Chen, Z., Lam, V., Han, J., Hassler, J., Finck, B. N., Davidson, N. O. & Kaufman, R. J. 2012. IRE1alpha-XBP1s induces PDI expression to increase MTP activity for hepatic VLDL assembly and lipid homeostasis. Cell Metab, 16, 473–86.

105. Wei, Y., Pattingre, S., Sinha, S., Bassik, M. & Levine, B. 2008a. JNK1-mediated phosphorylation of Bcl-2 regulates starvation-induced autophagy. Mol Cell, 30, 678–88.

106. Wei, Y., Sinha, S. & Levine, B. 2008b. Dual role of JNK1-mediated phosphorylation of Bcl-2 in autophagy and apoptosis regulation. Autophagy, 4, 949–51.

107. Welte, M. A. 2015. Expanding roles for lipid droplets. Curr Biol, 25, R470–81.

108. Willy, J. A., Young, S. K., Stevens, J. L., Masuoka, H. C. & Wek, R. C. 2015. CHOP links endoplasmic reticulum stress to NF-kappaB activation in the pathogenesis of nonalcoholic steatohepatitis. Mol Biol Cell, 26, 2190–204.

109. Wu, H., Ng, B. S. & Thibault, G. 2014. Endoplasmic reticulum stress response in yeast and humans. Biosci Rep, 34.

110. Yamaguchi, Y., Larkin, D., Lara-Lemus, R., Ramos-Castaneda, J., Liu, M. & Arvan, P. 2008. Endoplasmic reticulum (ER) chaperone regulation and survival of cells compensating for deficiency in the ER stress response kinase, PERK. J Biol Chem, 283, 17020–9.

111. Yamamoto, K., Takahara, K., Oyadomari, S., Okada, T., Sato, T., Harada, A. & Mori, K. 2010. Induction of liver steatosis and lipid droplet formation in ATF6alpha-knockout mice burdened with pharmacological endoplasmic reticulum stress. Mol Biol Cell, 21, 2975–86.

112. Younce, C. & Kolattukudy, P. 2012. MCP-1 induced protein promotes adipogenesis via oxidative stress, endoplasmic reticulum stress and autophagy. Cell Physiol Biochem, 30, 307–20.

113. Zhao, Y., Li, X., Cai, M. Y., Ma, K., Yang, J., Zhou, J., Fu, W., Wei, F. Z., Wang, L., Xie, D. & Zhu, W. G. 2013. XBP-1u suppresses autophagy by promoting the degradation of FoxO1 in cancer cells. Cell Res, 23, 491–507.

